# Epigenomic and chromosomal architectural reconfiguration in developing human frontal cortex and hippocampus

**DOI:** 10.1101/2022.10.07.511350

**Authors:** Matthew G. Heffel, Jingtian Zhou, Yi Zhang, Dong-Sung Lee, Kangcheng Hou, Oier Pastor Alonso, Kevin Abuhanna, Anthony D. Schmitt, Terence Li, Maximilian Haeussler, Brittney Wick, Martin Jinye Zhang, Fangming Xie, Ryan S. Ziffra, Eran A. Mukamel, Eleazar Eskin, Bogdan Pasaniuc, Joseph R. Ecker, Jesse Dixon, Tomasz J Nowakowski, Mercedes F. Paredes, Chongyuan Luo

**Affiliations:** Department of Human Genetics, University of California Los Angeles, Los Angeles, CA 90095, USA; Bioinformatics Interdepartmental Program, University of California Los Angeles, Los Angeles, CA 90095, USA; Genomic Analysis Laboratory, The Salk Institute for Biological Studies, La Jolla, CA 92037, USA; Bioinformatics and Systems Biology Program, University of California San Diego, La Jolla, CA 92093, USA; Department of Life Science, University of Seoul, Seoul 02504, Republic of Korea; Department of Pathology and Laboratory Medicine, David Geffen School of Medicine, University of California, Los Angeles, Los Angeles, CA 90095, USA; Department of Computational Medicine, David Geffen School of Medicine, University of California, Los Angeles, Los Angeles, CA 90095, USA; Department of Neurology, University of California San Francisco, San Francisco, CA, 94143, USA; Arima Genomics, Inc., 6354 Corte Del Abeto, Carlsbad, CA 92011, USA; Genomics Institute, University of California Santa Cruz, Santa Cruz, CA 95064, USA; Department of Epidemiology, Harvard T.H. Chan School of Public Health, Boston, MA 02115, USA; Program in Medical and Population Genetics, Broad Institute of MIT and Harvard, Cambridge, MA 02142, USA; Department of Cognitive Science, University of California, La Jolla, CA 92037, USA; Department of Neurological Surgery, University of California San Francisco, San Francisco, CA 94158, USA; Department of Anatomy, University of California San Francisco, San Francisco, CA 94158, USA; Department of Psychiatry and Behavioral Sciences, University of California San Francisco, San Francisco, CA 94158, USA; Eli and Edythe Broad Center for Regeneration Medicine and Stem Cell Research, University of California San Francisco, San Francisco, CA 94158, USA; Department of Computational Medicine, University of California Los Angeles, Los Angeles, CA 90095, USA; Howard Hughes Medical Institute, The Salk Institute for Biological Studies, La Jolla, CA 92037 USA; Peptide Biology Laboratory, The Salk Institute for Biological Studies, La Jolla, CA 92037, USA; Weill Institute for Neurosciences, University of California San Francisco, San Francisco, CA 94143, USA; Neuroscience Graduate Program, University of California San Francisco, San Francisco, CA 94143, USA; Developmental Stem Cell Biology, University of California San Francisco, San Francisco, CA 94143, USA

## Abstract

The human frontal cortex and hippocampus play critical roles in learning and cognition. We investigated the epigenomic and 3D chromatin conformational reorganization during the development of the frontal cortex and hippocampus, using more than 53,000 joint single-nucleus profiles of chromatin conformation and DNA methylation (sn-m3C-seq). The remodeling of DNA methylation predominantly occurs during late-gestational to early-infant development and is temporally separated from chromatin conformation dynamics. Neurons have a unique Domain-Dominant chromatin conformation that is different from the Compartment-Dominant conformation of glial cells and non-brain tissues. We reconstructed the regulatory programs of cell-type differentiation and found putatively causal common variants for schizophrenia strongly overlap with chromatin loop-connected, cell-type-specific regulatory regions. Our data demonstrate that single-cell 3D-regulome is an effective approach for dissecting neuropsychiatric risk loci.

## Main text

The adult human brain contains hundreds of cell types that display an extraordinary diversity of molecular, morphological, anatomic, and functional characteristics (*1–3*). Although the vast majority of cortical neurons are generated during the first and second trimesters, the highly distinct molecular signatures of cell types emerge between the third trimester and adolescence (*4–6*). Single-cell and bulk transcriptome analyses implicated dramatic gene expression remodeling in late prenatal and early postnatal development (*7, 8*). The pervasive transcriptome dynamics during human brain development is associated with genome-wide reconfiguration of DNA methylome and chromatin conformation (*9–12*). The brain-specific non-CG methylation (mCH, H=A, C or T) emerges in the human dorsal prefrontal cortex (PFC) during prenatal development in a cell type-specific pattern, with the average level of mCH increasing through adolescence (*9, 13*). This global reconfiguration of DNA methylation could profoundly shape neuronal development, for example, through the binding of mCH by MECP2, a gene implicated in neuropsychiatric disorders (*14–18*). In parallel with the accumulation of DNA methylation, chromatin architecture undergoes extensive remodeling in the first post-natal month of mouse brain development (*12*), which can potentially be mediated by neuronal activity-induced 3D genome rearrangement (*19*). The dynamic trajectory of DNA methylation and chromatin conformation changes have not been characterized with single-cell resolution in primary human brain tissues at the critical developmental stages of mid-gestation, late-gestation, and infancy. This study investigated the epigenomic dynamics in the developing human frontal cortex and hippocampus (HPC) using the multi-omic sn-m3C-seq approach to jointly profile chromatin conformation and DNA methylation in single nuclei (*20*).

We generated 29,691 sn-m3C-seq profiles (including 3,321 previously published (*20*)) from 13 developing and adult human frontal cortex samples, and 23,372 sn-m3C-seq profiles from 9 hippocampus samples (Fig. 1A, Fig. S1A-B, Table S1-2). We identified a total of 139 cell populations across all developmental stages by a fusion of three data modalities: mCH, CG methylation (mCG), and chromatin conformation (Fig. 1B, Table S3). These cell types are organized into 10 major groups (Fig. 1C). Excitatory neurons had distinct epigenomic types in the human PFC and HPC, which is consistent with their spatially separated *in situ* neurogenesis (Fig. 1D and Fig. S1C). By contrast, inhibitory neurons originating in the ventral portion of the embryonic brain, and non-neuronal cell types, are broadly shared between the two brain regions (Fig. 1D and Fig. S1C). Neuronal cell types, astrocytes, and oligodendrocyte progenitors were strongly separated by developmental stages based on their methylation and chromatin conformation patterns, whereas oligodendrocytes and other non-neural cell types showed similar epigenomic patterns across development (Fig. 1E). The developmental hierarchy of cortical and hippocampal cell types was reconstructed using hypo-methylation (either mCG or mCH) at cell-type marker genes, and computational integration of cells derived from different age groups using batch balanced k-nearest neighbor (BBKNN) based integration approaches (*21*) (Fig. 1F & Fig. S1D).

**Figure 1.**
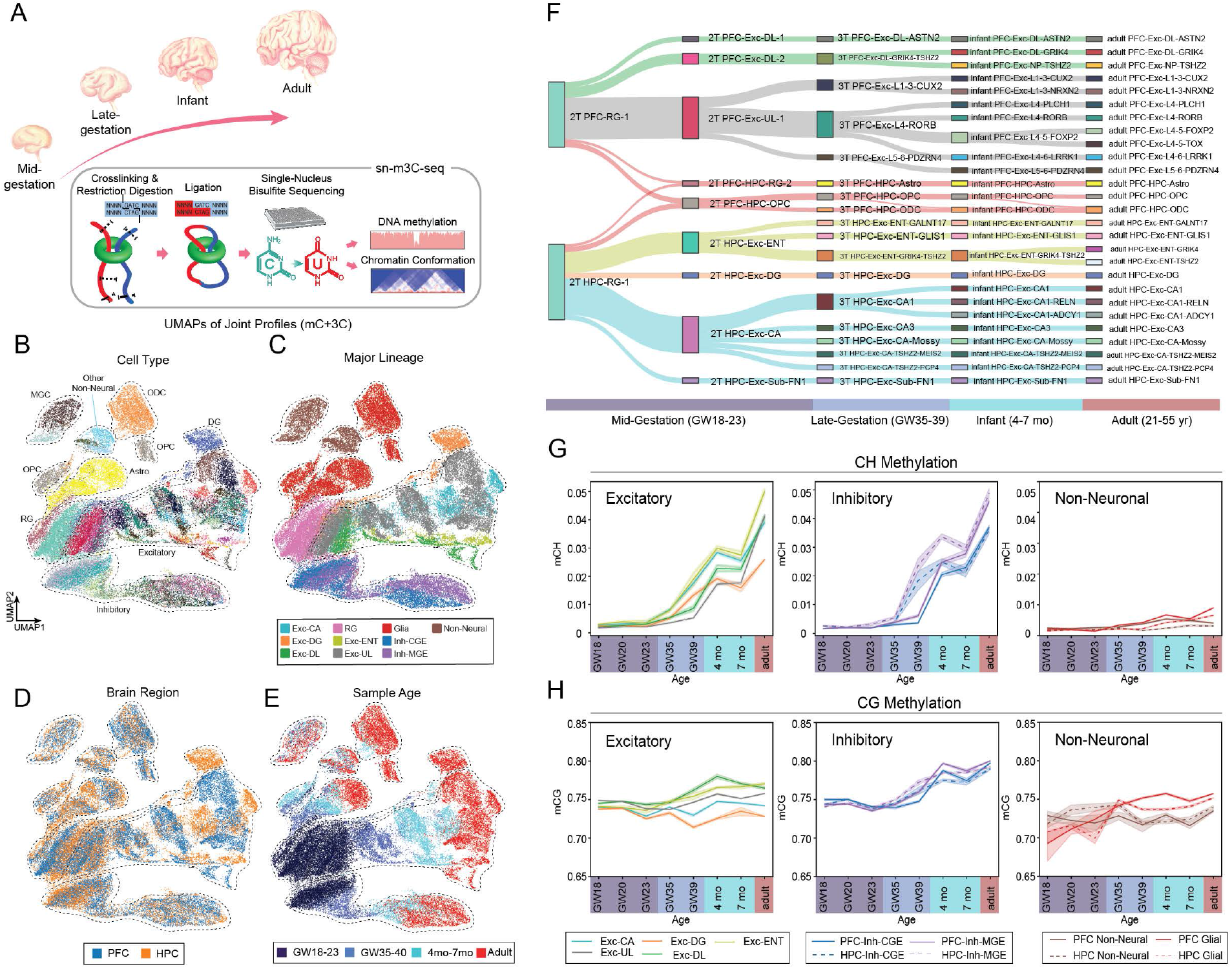
Profiling of epigenomic and chromatin conformation dynamics during human brain development using sn-m3C-seq. (A) Schematics of the study. (B-E) Dimensionality reduction using UMAP distinguishes cell types (B), major cell lineages (C), brain regions (D), and developmental stages (E). (F) Reconstructed developmental hierarchy of excitatory neurons and glial cells. (G-H) Dynamics of genome-wide non-CG methylation (G) and CG methylation (H) during human brain development.

The developmental increase of mCH in neuronal cells is an epigenomic hallmark of neuronal maturation (*9*). The accumulation of mCH in HPC excitatory and inhibitory neurons occurs earlier than that in PFC neurons (Fig. 1G), with HPC CA and inhibitory neurons containing significant amounts of mCH (>1% mean mCH/mCH) in two independent GW39 samples, whereas comparable mCH levels were not observed in PFC neurons until the infant stage (4 & 7 mo, Fig. 1G, Fig. S1E-G). Although genome-wide mCG levels only show moderate dynamics in neuronal populations (Fig. 1H), clustering of gene body mCG readily separates PFC neuronal populations into prenatal and postnatal groups (Fig. S1H), suggesting major reconfiguration of intragenic mCG between late-gestation and infant stages in PFC. A similar clustering analysis of HPC neuronal populations found that the remodeling of gene body mCG occurs in HPC between mid-gestation and late-gestation. The majority of neuronal populations in the late-gestational HPC, including CA1, CA3, dentate gyrus (DG), Mossy cell, and medial ganglionic eminence derived inhibitory neurons (Inh-MGE), were grouped with infant and adult cells rather than with mid-gestational cell types (Fig. S1I). To ask whether methylation remodeling in HPC also precedes that in PFC at individual loci, we selected genes whose mCG changes showed the strongest correlations with genome-wide mCG dynamics (see methods). All selected genes that show a gain of mCG during development underwent mCG remodeling in HPC ahead of PFC, as well as the vast majority (20/25) of genes showing developmental loss of mCG (Fig. S1J, K). Together, we found the remodeling of CH and CG methylome predominantly occurs between late gestation and infant stages, starting in the HPC and proceeding shortly thereafter in PFC.

Cell-type classification based on DNA methylation and chromatin conformation are largely concordant (Fig. S2A-D), with DNA methylation profiles providing a greater resolution for cell-type classification (Fig. S2D) (*20*). However, we found a notable exception in mid-gestational brains where a single neural progenitor radial glia (RG) population defined by DNA methylation signatures can be further divided using chromatin conformation signatures (Fig. S2E). Using chromatin conformation, we grouped RG cells into a neurogenic population RG-1 and a putative astrocyte progenitor population RG-2 (Fig. S2E) (*22*). This result was validated by an iterative classification of cells from mid-gestational brains, which found the gliogenic RG-2 more discretely defined by chromatin conformation than by DNA methylation (Fig. S2F). RG-1 can be further separated into an undifferentiated subpopulation, as well as cells that were primed for various excitatory cell-type trajectories (e.g. RG-CA, Fig. S2F). This observation led us to speculate that chromatin conformation dynamics precede the remodeling of DNA methylation during the differentiation of astrocytes. To test this hypothesis, we employed pseudotime analysis (*23*) to explore the temporal dynamics of chromatin conformation and DNA methylation by a more continuous time quantification than discrete donor ages. We computed pseudotime scores for RG-1, RG-2 and astrocyte cells using either gene body mCG or the interaction frequency of genomic bin pairs (Fig. S2G-H). In addition, to quantify chromatin conformation at individual loci, we devised “3C (Chromatin Conformation Capture) gene score” representing the sum of intragenic chromatin contact frequency. The results suggest little mCG dynamics in RG-1 and RG-2 populations, and substantial remodeling of mCG in differentiated astrocytes (Fig. S2G). In contrast, the reconfiguration of chromatin interactions was more continuous across the differentiation of RG to astrocytes (Fig. S2H), which resulted in a drastically different distribution of pseudotime scores computed from mCG or chromatin interactions (Fig. S2I). The comparison of pseudotime scores of the two data modalities in the same cells further revealed a dramatic separation of the temporal dynamics of mCG and chromatin conformation (Fig. S2J-K). The differentiation of RG to astrocytes can be divided into a stage of rapid chromatin conformation remodeling in RG-1 and RG-2 that predominantly occurs during mid-gestation, followed by a notably protracted maturation of CG methylome that extends into the adult brain (Fig. S2I-J). Consistent with genome-wide pseudotime patterns, gene-specific analyses found the remodeling of 3C gene score generally occurs in RG-1 and RG-2 populations and precedes mCG dynamics in differentiated astrocytes (Fig. S2L-O). The protracted mCG dynamics in astrocytes were also supported by a chromatin loop analysis that identified 5,904 differential loops overlapping with 1,432 genes during astrocyte differentiation (Fig. S2P-Q, Table S4). While the majority of chromatin loops were reconfigured in mid-gestation when RG-1 differentiates to RG-2 (Fig. S2P), the gain or loss of gene body mCG did not occur until the infant or adult stages (Fig. S2Q). Consistent with the transition from RG-1 to glial progenitor RG-2 and further to astrocytes, the genes overlapping with gained chromatin loops are enriched in gene ontology terms such as “axon target recognition” (Fig. S2R), whereas genes overlapping with lost chromatin loops are enriched in terms such as “glial cell fate specification” (Fig. S2S). The differentiation of astrocytes is also associated with rearrangements of chromatin domain boundaries. We identified 684 differential domain boundaries during astrocyte differentiation (Table S5). For example, the developmental strengthening of a domain overlapping with astrocyte marker gene SLC1A2 is associated with considerable loss of gene body mCG in the infant stage (Fig. S3A). In contrast, the early developmental gene SOX11 locates at the boundary of two domains in RG-1. The boundary is diminished during development and becomes undetectable in late gestation (Fig. S3B).

We asked whether the temporal separation of DNA methylation and chromatin conformation dynamics is shared by other cell-type differentiation trajectories focusing on MGE-derived inhibitory neurons that are shared between PFC and HPC (Fig. 2A-B). We calculated pseudotime scores using both gene body mCG and the interaction frequency of genomic bin pairs and observed a dramatic difference in the distribution of pseudotime scores for methylation and chromatin conformation in inhibitory maturation (Fig. 2C). Unlike the protracted methylation remodeling observed in astrocyte differentiation (Fig. S2J), the pseudotime of mCG exhibited a bimodal distribution during the maturation of MGE-derived inhibitory neurons, suggesting a rapid transition between the methylation states of developing and mature cells (Fig. 2C). Indeed, the vast majority of mCG pseudotime range was traversed by a single cell type, Inh-MGE, and predominantly during late gestation (Fig. 2D-E). Conversely, the pseudotime for 3C gene score was more uniformly spaced in a trimodal distribution, presenting two distinct transitory events (Fig. 2C). Chromatin conformation pseudotime traverses the least amount of pseudotime distance in Inh-MGE with the majority of distance traveled before, in Inh-eMGE during mid-gestation (stage 1 in Fig. 2D-E), or after, in Inh-MGE-ERBB4 during infant to adult stages (stage 2 in Fig. 2D-E). Further, in agreement with our earlier analyses, we saw in the pseudotime scores that MGE-derived inhibitory neurons located in HPC matured more quickly than those located in PFC, with the difference being more pronounced for mCG than for chromatin conformation (Fig. 2F-G). Consistent with the two stages of chromatin conformation dynamics (Fig. 2D-E), we found loci primarily lost 3C gene score during mid-gestation (Stage 1, Fig. 2H) and predominantly gained 3C gene score during infant and adult stages (Stage 2, Fig. 2I). A parallel analysis on a caudal ganglionic eminence (CGE) derived inhibitory neuron trajectory found similar results (Fig. S4), but with a more uniform distribution in chromatin conformation pseudotime than in the MGE-derived analyzed trajectory (Fig. S4B-C).

**Figure 2.**
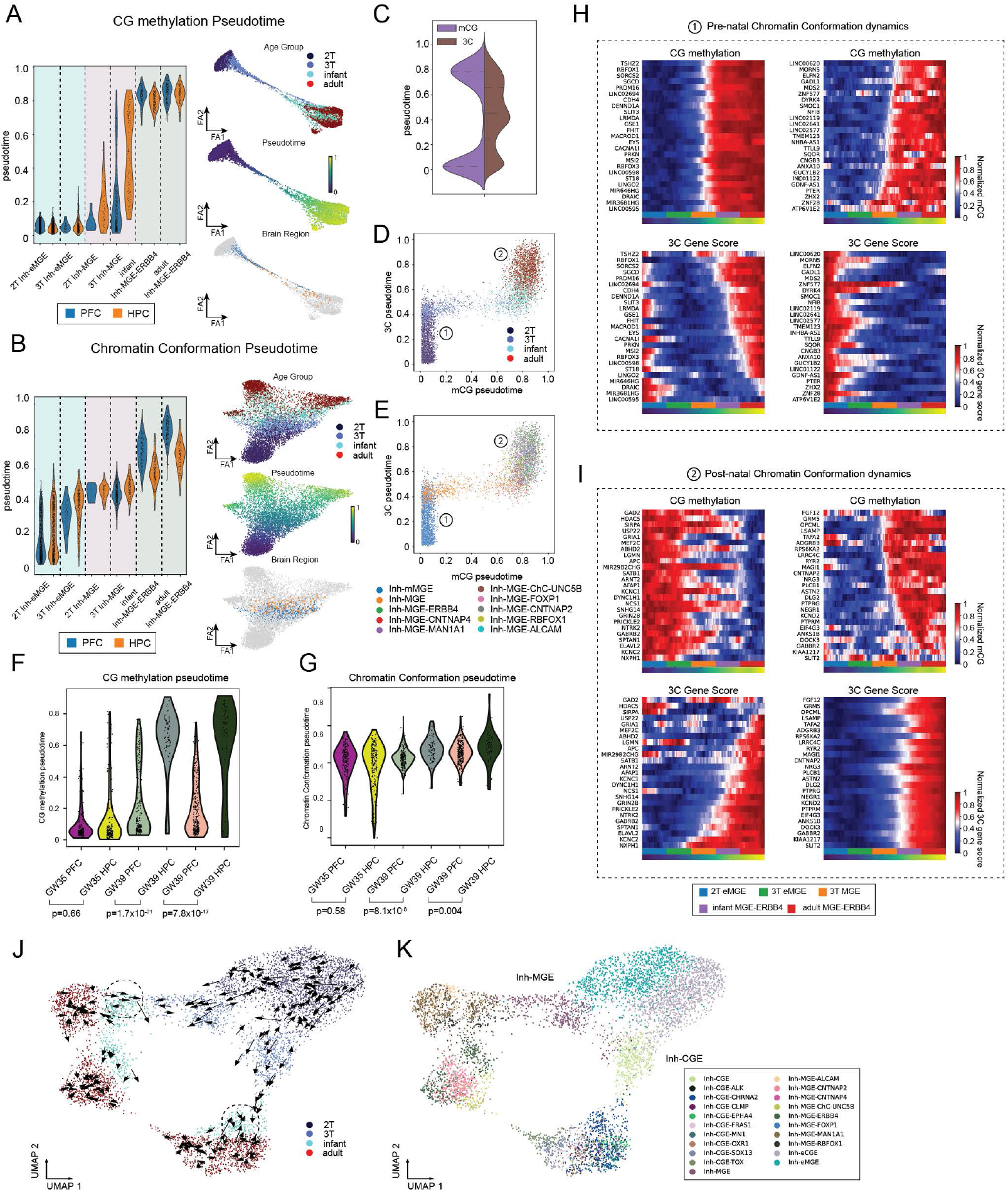
Temporal ordering of DNA methylation and chromatin conformation reconfiguration during the maturation of MGE-derived inhibitory neurons. (A-B) Distribution of pseudotime scores computed from gene body mCG (A) or interaction frequency of genomic bin pairs (B) in MGE-derived inhibitory neurons. (C) Distinct distributions of pseudotime scores computed from mCG or chromatin conformation. (D-E) Direct comparison of pseudotime scores computed from mCG or chromatin conformation in individual cells, labeled by developmental age groups (D) and cell types (E). (F-G) Comparison of mCG (F) or chromatin conformation (G) pseudotime score distributions for MGE-derived inhibitory neurons located in PFC and HPC, using three late-gestational brain samples from which both PFC and HPC were analyzed. (H) mCG and 3C gene scores changes at loci associated with pre-natal chromatin conformation dynamics. (I) mCG and 3C gene scores changes at loci associated with pot-natal chromatin conformation dynamics. (J-K) 3D chromatin potential analysis of inhibitory neuron differentiation. Dimensionality reduction plots were labeled for developmental stages (J) or cell types (K). Reverse flows from adult and infant stages to late-gestation were highlighted.

We devised a “3D chromatin potential” analysis based on the “chromatin potential” approach originally developed to temporally order chromatin accessibility and gene expression changes (*24*), to ask whether the genome-wide chromosome conformation at a developmental stage could indicate its future methylation landscape. For each cell, we identified 5 cells whose methylation profiles are the most compatible with its 3C profile (mCG neighbors of a 3C profile). Each arrow represents the distance between the true mCG profile of a single cell to its mCG neighborhood, best predicted by its chromatin conformation state (Fig. 2J-K). During the differentiation of MGE-derived inhibitory cells, 3D chromatin potential initially flows from mid-gestation to late-gestation (Fig. 2J-K), which corresponds to the chromatin conformation rearrangement in mid-gestation and a lag of mCG remodeling (stage 1 in Fig. 2D-E). Following the abrupt mCG change during late-gestation, significant reversal flows of 3D chromatin potential from the adult and infant stage to late-gestation were observed and were likely driven by the chromatin conformation remodeling during infant and adult development (stage 2 in Fig. 2D-E).

The prediction of cell-type-specific gene expression using mCH in the adult brain is well-established (*25, 26*). Here we have extended the approach to the developing brain taking advantage of the inverse correlation between gene expression and mCG or mCH (Fig. S5A-D). We used single-molecule fluorescent *in situ* hybridization to investigate the RNA expression patterns of cell-type markers identified by the methylation analyses. TLL1, a gene that shows reduced gene body mCG in granule cell layer (GCL) neurons, was localized to the GCL in the hippocampus in the third trimester (GW 30) (Fig. S5B, E). There were overlaps with RBFOX3, a molecular marker for mature neurons, and PROX1, a transcription factor found in granule neurons of the hippocampus (Fig. S5E). TRPS1 mRNA was expressed in excitatory (GAD1-negative) cells in the hilus and CA3 regions, supporting it as expressed in mossy cells and CA3 pyramidal neurons in the third trimester (Fig. S5C,F). Lastly, we identified an increased 3C gene score at the LRIG1 locus in the putative astrocyte progenitor population RG-2 and reduced mCG in astrocytes (Fig. S2N-O and Fig. S5D). We found a significant fraction (40%) of cells expressing a canonical astrocyte marker ALDH1L1 also express LRIG1 (Fig. S5G-H), supporting the dynamic expression of LRIG1 during astrocyte differentiation.

Chromatin conformation capture techniques produce snapshots of 3D genome architecture at multiple scales, including A/B compartments and more local features such as chromatin domains and loops (*27*). While A/B compartments are detected through long-range interactions (e.g. > 10 Mb distance between interacting loci), chromatin domains and loops are primarily detected by median-range interaction with less than 5 Mb distance. We clustered single-cell 3C profiles by the distribution of the distance between interacting loci using k-means clustering and found single brain cells range from mainly containing median-range interaction (Cluster 1) to primarily containing long-range interactions (Cluster 10) (Fig. 3A-B, Fig. S6A-B). Strikingly, neuronal cell types are strongly enriched in Clusters 1-6, dominated by median-range interactions, whereas glial and non-neural cell types are enriched in Clusters 8-10 that are dominated by long-range interactions (Fig. 3C and Fig. S6C). We have developed thresholds to categorize the global chromatin conformation of each cell into Domain Dominant (DD), Compartment Dominant (CD) as well as Intermediate (INT) (Fig. 3D and Fig. S6D-E). We analyzed a published bulk Hi-C dataset generated from primary human tissues and found bulk chromatin conformation profiles from all 10 tissues show CD signature (*28*), suggesting that DD is a global chromatin conformation specific to neuronal cells (Fig. 3D). Interestingly, although Hi-C profiles generated from bulk human cortical and hippocampal tissues show a greater fraction of median-range interaction than other somatic tissues, they were nevertheless classified as CD-type samples likely due to the abundant non-neuronal cells in the analyzed tissue (Fig. 3D). This observation highlights the advantage of single-cell 3C profiling in discerning cell-type specific chromatin conformation profile in heterogeneity tissues. The differentiation of neurons and astrocytes involved distinct remodelings of the global chromatin conformation (Fig. 3E-G, Fig. S6F-J). The neural progenitor RG-1 population is depleted of CD conformation but is not enriched in either DD or INT conformations. The chromatin conformation was rapidly remodeled in progenitors committed to producing upper-layer excitatory neurons (RG-UL) and young neurons in the mid-gestational brain (2T-Exc-UL) and showed a comparable enrichment in DD conformation as in adult neurons (Fig. 3E,G). The differentiation of astrocytes involved a transition to CD conformation, which was completed during late-gestation (Fig. 3F-G).

**Figure 3.**
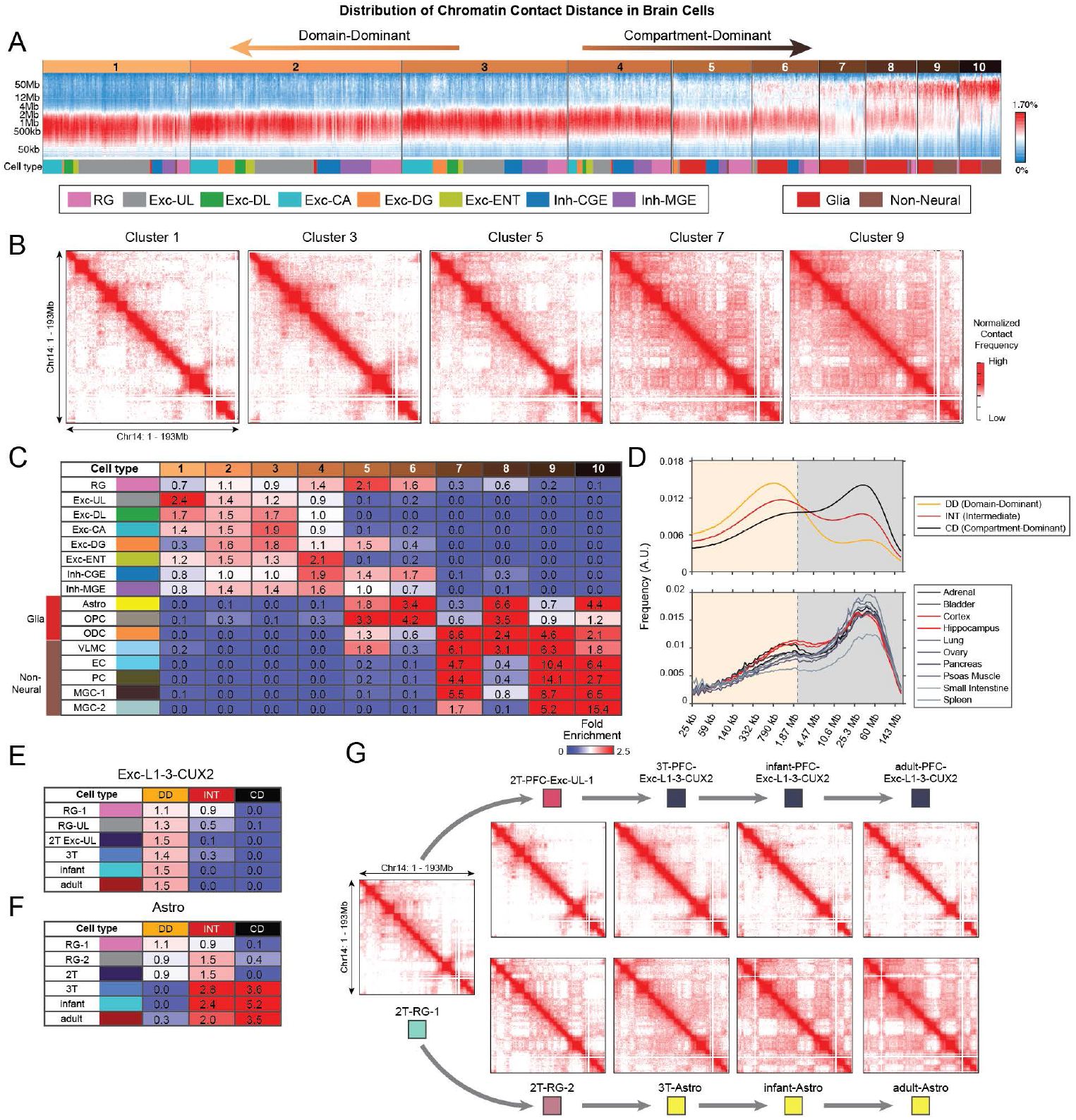
Remodeling of global chromatin conformation during human brain development. (A) K-means clustering analysis groups single-cell 3C profiles by the distribution of the distance between interacting loci. (B) Merged chromatin interaction profiles of odd clusters identified in (A). (C) Cell-type specific enrichments of clusters identified in (A). (D) Comparison of Domain-Dominant (DD), Compartment-Dominant (CD), and Intermediate (INT) chromatin conformation found in single brain cells to bulk Hi-C profiles of diverse human tissues. (E-F) Remodeling of global chromatin conformation during the differentiation of upper-layer excitatory neurons (Exc-L1-3-CUX2) (E) and astrocytes (Astro) (F) from the common RG-1 progenitor. (G) Merged chromatin interaction profiles of developing cell populations across the differentiation of upper-layer excitatory and astrocytes.

Differentially methylated regions are a reliable marker of dynamic regulatory activity, with loss of methylation indicating an increase in regulatory activity and gaining of methylation associated with repression (*29, 30*). We investigated the global regulatory dynamics of human cortical and hippocampal development by identifying over 2.5 million differentially methylated regions (DMRs) across all cell types and developmental stages (Fig. 4A, Fig. S7A), followed by the analysis of transcription factor (TF) binding motif enrichment (Fig. 4B, Fig. S7B). Excitatory and inhibitory cells from the prenatal brain and glial cells from all stages share a large amount of DMRs that are enriched in binding motifs for EMX, LHX2, SOX, and DLX TFs (cluster 1 in Fig. 4A). This is consistent with earlier analyses in Fig. S1H-I showing that the epigenomic difference between excitatory and inhibitory neurons are moderate in the pre-natal brain and becomes much more pronounced in infant and adult brains (Fig. S1H-I). While all excitatory populations share a group of pan-excitatory DMRs that are enriched in the Neurogenin and RFX binding motifs (cluster 3), most excitatory subtypes such as HPC-CA, PFC-DL (Deep-Layer), PFC-UL (Upper-Layer), and Mossy cells are associated with their sub-population specific DMRs (clusters 4-7). Consistent with a higher expression of LHX2 in the developing hippocampus (*31*), Exc-CA and Exc-DG show a stronger enrichment of LHX2 motifs than in cortical excitatory cells (Fig. 4B, Fig. S7B).

**Figure 4.**
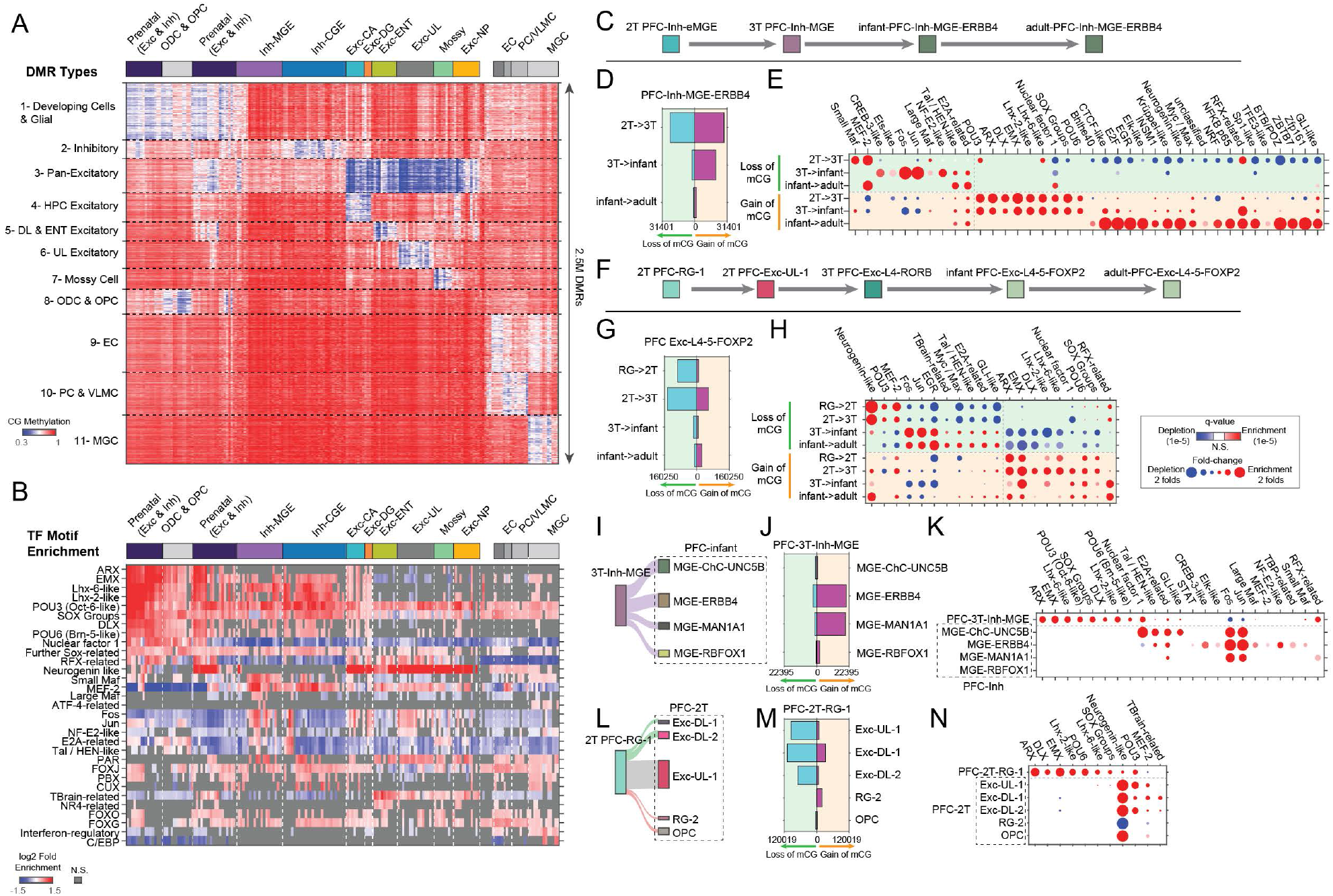
Regulatory dynamics during brain cell type differentiation. (A) K-means clustering of differentially methylation regions (DMRs) reveals specificities for cell lineages and developmental stages. (B) Transcription binding motif enrichment analysis of DMRs. (C) Schematics of the maturation of MGE-derived ERBB4-expressing inhibitory neurons (Inh-MGE-ERBB4). (D) Numbers of trajectory-DMRs identified for Inh-MGE-ERBB4 maturation between adjacent developmental stages. (E) Enriched transcription factor binding motifs in trajectory-DMRs for the maturation of Inh-MGE-ERBB4 neurons. (F) Schematics of the differentiation of FOXP2-expressing excitatory neurons (Exc-L4-5-FOXP2). (G) Numbers of trajectory-DMRs identified throughout the differentiation of Exc-L4-5-FOXP2 neurons. (H) Transcription factor binding motif enrichments in trajectory-DMRs for the differentiation of Exc-L4-5-FOXP2. (I) Schematics of the specification of MGE-derived inhibitory neuron types. (J) Numbers of branch-DMRs found during the specification of MGE-derived inhibitory neuron types. (K) Transcription factor binding motif enrichments in branch-DMRs for MGE-derived inhibitory neuron types. (L) Schematics of the specification of RG-1 derived cell types. (M) Numbers of branch-DMRs found during RG-1 differentiation. (N) Transcription factor binding motif enrichments in branch-DMRs associated with RG-1 differentiation.

Although epigenomic analyses of adult mammalian brains have identified lineage-specific TF activities (*26, 32, 33*), the developmental dataset generated in this study allows us to infer the temporal sequence of TF activity. We have identified dynamic DMRs across the stages of cell-type specification (trajectory-DMRs, Fig. 4C-H, Fig. S8-9) and DMRs that distinguish daughter cell populations derived from a common mother cell type (branch-DMRs, Fig. 4I-N, Fig. S10-11). A distinct wave of repression of regulatory elements (gain of mCG DMRs) during the late-gestation was found during the differentiation of cortical inhibitory neurons (Fig. 4C-D and Fig. S9). Consistent with the early maturation of hippocampal neurons, the wave of repression for regulatory elements was found during mid-gestation for hippocampal inhibitory neurons (Fig. S9C,E,F,H). The differentiation of RG to diverse excitatory neurons during mid-gestation is associated with pervasive activation of regulatory elements, as shown by the numerous DMRs that lose mCG (Fig. 4F-G and Fig. S8B-C,F,G-I). Consistent with the protracted maturation of astrocyte methylome (Fig. S2I-J), the maturation of astrocytes between infant and adult brains is associated with a loss of mCG at 57,783 regions, far exceeding the scale of mCG remodeling in earlier stages of astrocyte differentiation (Fig. S8D). Using TF binding motif analysis, we found the regulatory landscape of both excitatory and inhibitory neurons is shaped by the sequential action of lineage-specific and activity-dependent TFs. Regulatory elements that become activated (loss of mCG) in the mid-gestation are enriched in the binding motifs of lineage-specific TFs such as Maf and MEF-2 for inhibitory cells (Fig. 4E), or Neurogenin, MEF-2 and POU3 for excitatory neurons (Fig. 4H). Following lineage specification, the binding motif of activity-dependent TFs (FOS, JUN, EGR1, CREB) is strongly enriched in regulatory elements activated in late-gestation to infant stages in both excitatory and inhibitory populations (Fig. 4E,H) (*34*). This result suggests late-gestational to early-infant development as a key stage during which the epigenome is shaped by neuronal activity. The analysis of branch-DMRs supported the wave of repression for regulatory elements in diverse subtypes of inhibitory neurons during late gestation (Fig. 4I-K). Furthermore, the analysis of RG2 differentiation supports the gliogenic characteristic of this progenitor pool as the binding motif of Neurogenic TFs is strongly depleted in regions losing mCG in RG-2 (Fig. 4L-N).

Using DMRs and chromatin loops identified from adult and developmental cell types, we systematically localized the heritability signals of neuropsychiatric disorders across developmental stages and cell populations. The polygenic heritability enrichment of annotations defined by DMR and/or chromatin loops was quantified for each cell type using stratified linkage disequilibrium score regression (S-LDSC) (*35*) (Figure S12-13). We found significantly greater enrichment of heritability in loop-connected DMRs (DMRs that overlap with a chromatin loop) than in all DMRs (Fig. 5A and Fig. S14A-F, p = 1.8×10^−49^ via paired t-test), supporting the utility of chromatin loops in locating potential causal variants. The specificity of our analysis was validated by the specific enrichment of the heritability of Alzheimer’s disease in microglia and the absence of strong enrichment for height heritability across analyzed cell types (Fig. S12). We also overlapped fine-mapped putative causal loci of schizophrenia (*36*) to DMRs and loop-connected DMRs (190 independent loci containing 569 high-confidence putative causal SNPs with posterior inclusion probability (PIP) > 0.1, Table S6). Out of 190 schizophrenia fine-mapped loci, 111 and 81 loci contain at least one putative causal SNPs that overlap with a DMR or loop-connected DMR, respectively (Fig. 5B). We found a strong correlation between the odds ratio of overlapping with a putative causal SNP, and the enrichment of polygenic heritability across cell types (Fig. 5C, Spearman’s correlation = 0.74, p = 8.6×10^−31^). As an example, we showcase rs500102 (PIP=0.27), a putative causal variant for schizophrenia, that overlaps with a loop-connected DMR in L4-5 excitatory neurons (Fig. 5D). The variant is also a fine-mapped eQTL of RORB detected in the brain tissue by GTEx studies (Table S6) (*37*). The region where rs500102 is localized is connected by a loop domain to RORB promoter, specifically in L4-5 excitatory neurons (Fig. 5D). The loop domain is associated with cell-type-specific reduction of mCG in RORB gene body as well as in the region surrounding rs500102 (Fig. 5D). The example demonstrates the utility of single-cell multi-omic profiles to generate mechanistic hypotheses regarding the function of GWAS-associates variants.

**Figure 5.**
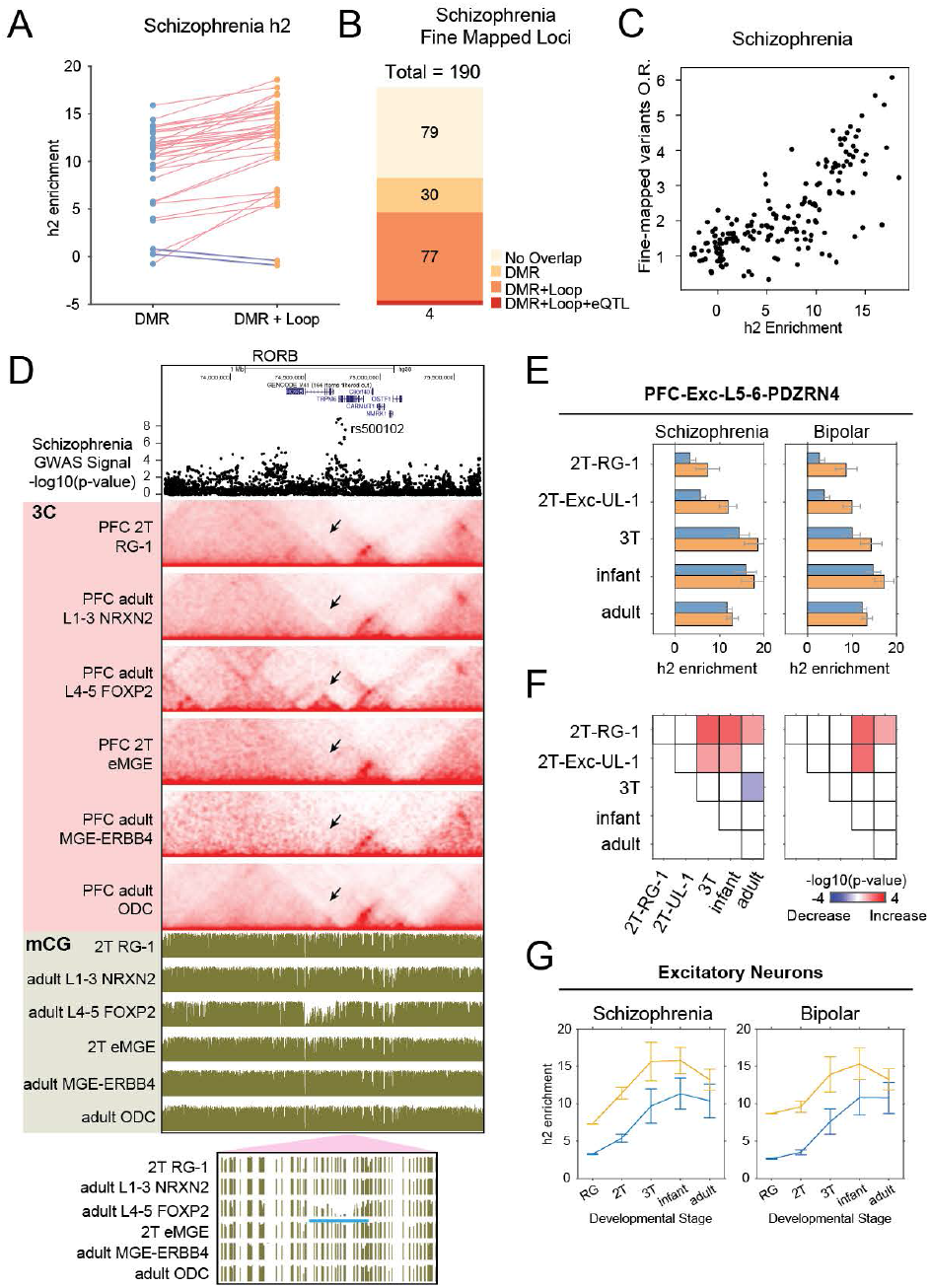
Localizing the heritability signals of neuropsychiatric disorders across developmental stages and cell populations. (A) The enrichment of schizophrenia polygenic heritability in DMRs and loop-connected DMRs. (B) Numbers of schizophrenia-associated loci containing at least one fine-mapped variant that overlaps with DMRs or loop-connected DMRs. (C) Correlation between the enrichments of polygenic heritability and fine-mapped schizophrenia variants. (D) The genomic region overlapping with a putative causal variant for schizophrenia rs500102 is connected to the RORB promoter through a cell-type-specific loop domain. (E) Enrichment of polygenic heritability for schizophrenia and bipolar disorder in PDZRN4-expressing Layer 5-6 excitatory neurons across developmental stages. (F) Statistical significance of differential heritability enrichment between development stages. P-values were computed two-sides t-tests. Red and blue colors show developmentally increase or decrease of heritability enrichment, respectively. (G) Meta-analysis of heritability enrichment for schizophrenia and bipolar disorder in excitatory neuron populations.

Next, we assessed the developmental dynamics of enrichment for neuropsychiatric disorder heritability in various neuronal populations (Fig. 5E-F and Fig. S14G-L). For schizophrenia and bipolar disorder, the enrichment of polygenic heritability increases from neuroprogenitor (RG-1) to early post-mitotic neurons (e.g. 2T Exc-UL-1) and further to post-mitotic neurons in late-gestational brains for both excitatory (Fig. 5E-F, Fig. S14H-J) and inhibitory populations (Fig. S14K-L). We also found a trend of decreased heritability enrichment in adult neurons for schizophrenia and bipolar disorder, although the decreases are not statistically significant except for in a L5-6 excitatory population (Fig. 5E-F). Using meta-analyses of all excitatory (Fig. 5G, Fig. S14M) or inhibitory populations (Fig. S14N), we found a consistent developmental increase of enrichment for schizophrenia and bipolar disorder between neuroprogenitors and neurons in infant brains, followed by a decrease in the adult brain. Taken together, our results suggest that the genetic risk of schizophrenia and bipolar disorder more strongly affects post-mitotic neurons than the neuroprogenitor population in developing human brains.

Genome-wide rearrangements of DNA methylome is crucial for the normal development of mammalian brains. The disruption of the *de novo* “writer” of DNA methylation DNMT3A (*38, 39*), or the “reader” of DNA methylation MECP2, cause molecular to behavioral alterations in animal models (*39*) and lead to human neuropsychiatric conditions such as the DNMT3A-associated Tatton Brown Rahman Syndrome, or the MECP2-associated Rett Syndrome (*40, 41*). Using single-cell multi-omic profiling, our study found the remodeling of neuronal methylome predominantly occurs during late-gestational and early-post-natal development, suggesting the human brain is particularly vulnerable to genetic and environmental perturbations that impact DNA methylation functions in these developmental stages. In addition to the exceptional abundance of non-CG methylation, we found another layer of unique epigenomic regulation in neurons, that is, the unusually strong chromatin domain strength that is different from glial cells or non-brain tissues. The finding raises interesting questions regarding whether cohesin-dependent enhancer-promoter loops are regulated differently in neurons than in non-neuronal cell types (*42*). The single-cell multi-omic dataset generated by sn-m3C-seq provides cell-type-specific functional annotations (i.e. DMRs and chromatin loops) to more than half of fine-mapped schizophrenia-associated loci, highlighting the application of sn-m3C-seq profiles in dissecting the developmental context and molecular mechanism of non-coding variants associated with neuropsychiatric disorders.

## Data Access

Datasets generated by this study can be accessed interactively through https://brain-epigenome.cells.ucsc.edu/. Processed chromatin conformation data for all samples, and processed single-cell DNA methylation data for unrestricted access samples can be downloaded at NCBI GEO accession GSE213950. Raw sequencing reads for both controlled and unrestricted access samples can be downloaded from NeMO Archive.

## Code Availability

Codes for the demultiplexing of sn-m3C-seq fastq files are available at https://github.com/luogenomics/demultiplexing. Modified TAURUS-MH for mapping of sn-m3C-seq data is available at https://github.com/luogenomics/Taurus-MH. Codes for the generation and imputation of methylation features is available at https://github.com/luogenomics/snm3Cseq_feature_processing.

## Author contributions

C.L., T.J.N, and M.F.P. conceived the project. O.P.A., R.Z., and M.F.P. collected and processed the tissue specimens and dissected the samples. Y.Z., A.D.S., and C.L. optimized the sn-m3C-seq method. Y.Z., K.A., and C.L. generated the data. M.G.H., J.Z., D-S.L., K.H., J.D., T.L., M.J.Z., and F.X. analyzed the data. C.L., M.F.P., T.J.N, J.D., J.R.E., B.P., E.E., and E.A.M. managed the project. C.L., M.G.H., and K.H. draft the manuscript. K.A., M.H., and B.W. developed the data browser. All authors edited the manuscript.

## Acknowledgment

We thank Drs. Noah Zaitlen (UCLA), Michael Gandal (UCLA), Jonathan Flint (UCLA), and Alkes Price (Harvard) for discussions; Byron Ashley for illustration. This work is supported by NIMH awards R01MH125252 and U01MH130995 to C.L; NIMH award U01MH121282 and NHGRI award R01HG010634 to J.R.E; NIMH award U01MH116438, NINDS award R01NS123263 and Simons Foundation grant SF810018 to T.J.N. P01 NS083513, Roberta and Oscar Gregory Endowment in Stroke and Brain Research, Chan Zuckerberg Biohub, and 1DP2NS122550-01 to M.F.P. NIMH award R01MH115676 to B.P. NHGRI award 2U24HG002371-23 to M.H. Institute of Information & Communications Technology Planning & Evaluation (IITP) of Korea award IITP2022-0-006240101005, 2021 Research Fund of the University of Seoul (Grant No. 202104151024), National Research Foundation (NRF) of Korea awards NRF2021R1C1C1006798, NRF2021M3H9A209722712, and NRF2022M3H9A108101112 to D.L. J.R.E is an investigator of the Howard Hughes Medical Institute.

## Competing interests

J.R.E. serves on the scientific advisory board of Zymo Research Inc.

## Supplementary Materials

### Materials and Methods

Supplementary Table 1. Human Brain Specimen used in this study.

Supplementary Table 2. sn-m3C-seq metadata

Supplementary Table 3. Cell Type Annotation

Supplementary Table 4. Differential chromatin loops.

Supplementary Table 5. Differential chromatin domain boundaries.

Supplementary Table 6: Results for overlapping schizophrenia fine-mapped high confidence putative causal loci (PIP > 10%) with DMR / loop / eQTL.

Supplementary Table 7. Sequences of 384-plex RP-H primers

## Supplementary Figure Legends

**Supplementary Figure 1. Related to Figure 1.**
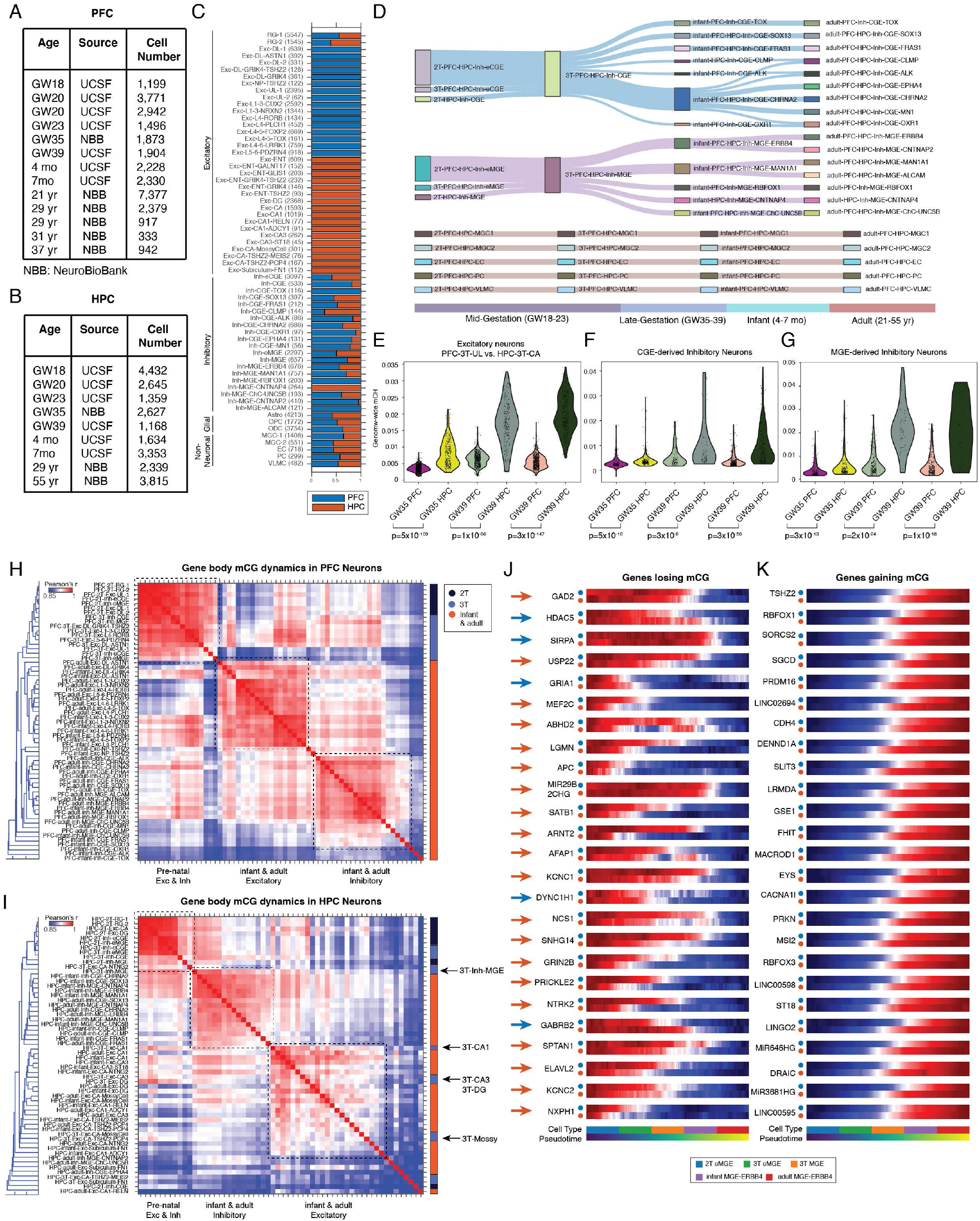
(A-B) Numbers of sn-m3C-seq profiles generated from each specimen. (C) Brain regional specificity of identified cell types. (D) Reconstructed developmental hierarchy of inhibitory neurons and non-neuronal cells. (E-G) Comparison of genome-wide mCH level in cortical and hippocampal excitatory neurons (E), CGE-derived inhibitory neurons (F), and MGE-derived inhibitory neurons (G) in the late-gestational samples. (H) Correlation matrix of PFC neuronal populations computed with gene body mCG. (I) Correlation matrix of HPC neuronal populations computed with gene body mCG. (J-K) Comparison of the timing of mCG remodeing in PFC and HPC for genes showing developmentally loss of mCG (J) and gain of mCG (K).

**Supplementary Figure 2. Related to Figure 2.**
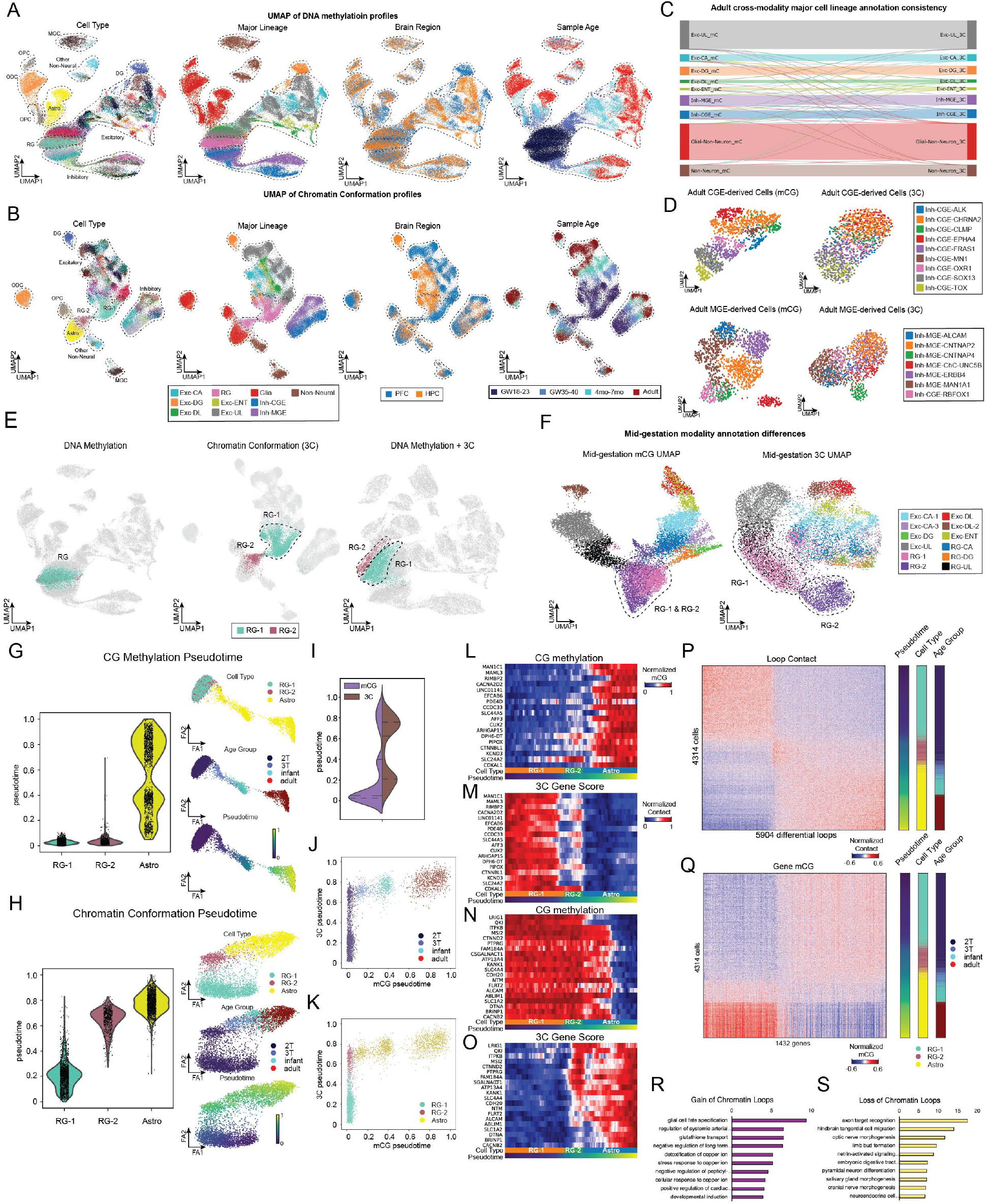
(A) Methylation dimensionality reduction using UMAP distinguishes, from left to right, cell types, major cell lineages, brain regions, and sample age groups. (B) Chromatin conformation dimensionality reduction using UMAP distinguishes, from left to right, cell types, major cell lineages, brain regions, and sample age groups. (C) Riverplot to show extensive consistency in adult major cell lineage annotation between DNA methylation and chromatin conformation modalities. (D) Dimensionality reduction using UMAP shows resolution difference in cell-type classification using DNA methylation and chromatin conformation modalities in adult inhibitory neurons; CGE-derived (top), MGE-derived (bottom). (E) Showing distinction of RG-1 and RG-2 populations in methylation space (left), chromatin conformation space (middle), and joint dimensionality reduction space (right). (F) Dimensionality reduction of z-scored mCG feature matrix for cells from mid-gestational brains (left). Chromatin conformation dimensionality reduction of cells from mid-gestational brains (right). (G-H) Distribution of pseudotime scores computed from gene body mCG (G) or chromatin conformation (H) in neural progenitor RG-1, astrocyte progenitor RG-2 and astrocyte populations. (I) Distinct distributions of pseudotime scores computed from mCG or chromatin conformation. (J-K) Direct comparison of pseudotime scores computed from mCG or chromatin conformation labeled by sample age group (J) and cell types (K). (L-M) Highest ranking genes by mCG correlation to the pseudotime displayed as mCG gene body (L) and 3C gene score (M). (N-O) Highest ranking genes by mCG anticorrelation to the pseudotime displayed as mCG gene body (N) and 3C gene score (O). (P) Chromatin contact frequency at differential chromatin loops identified during astrocyte differentiation. (Q) Intragenic mCG for genes overlapping with differential chromatin loops identified during astrocyte differentiation. (R-S) Gene ontology term enrichments for genes overlapping with gained chromatin loops (R) and lost chromatin loops (S).

**Supplementary Figure 3. Related to Figure 2.**
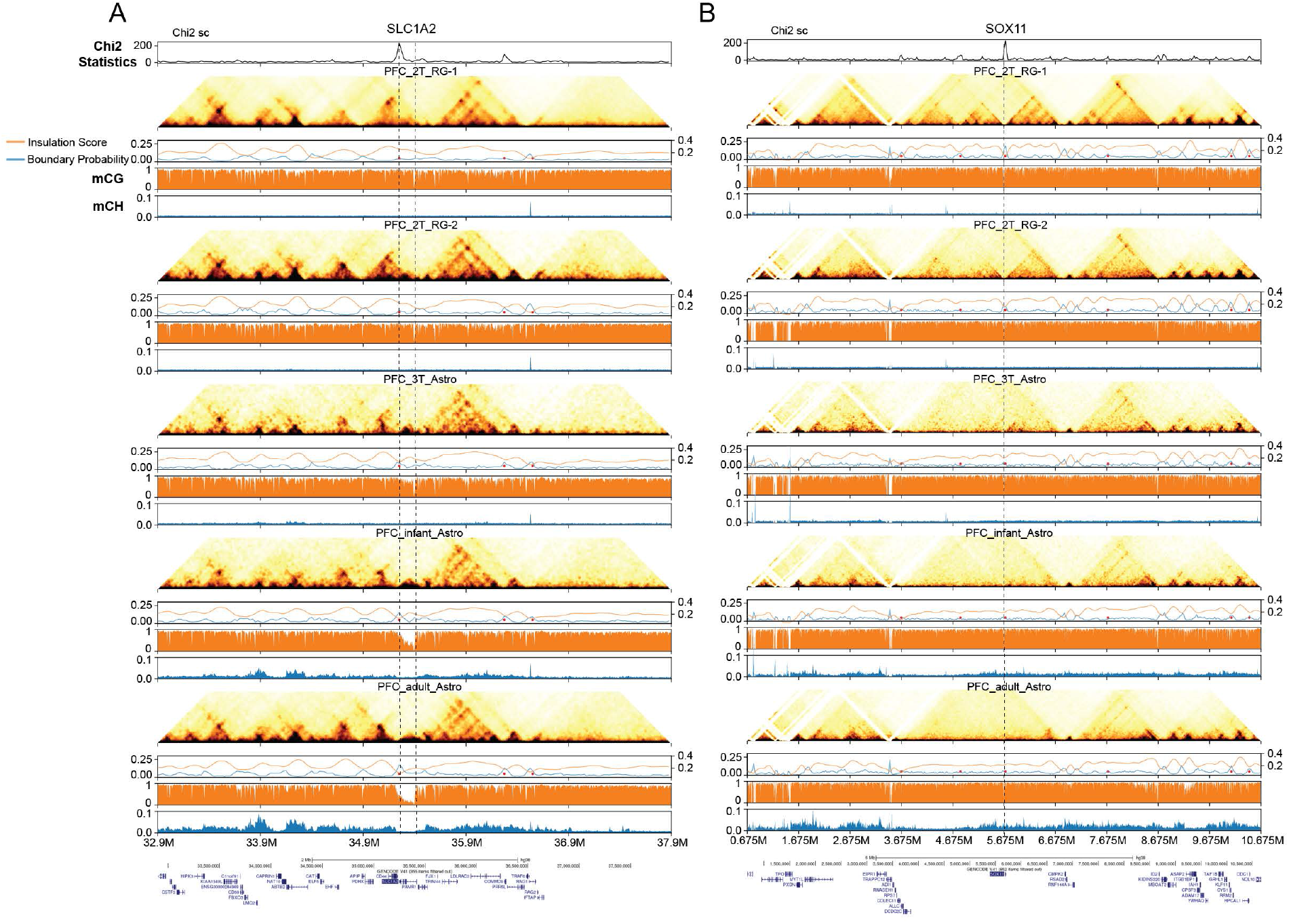
Differential chromatin domain boundaries were identified during the differentiation of astrocytes at SLC1A2 (A) and SOX11 (B) loci.

**Supplementary Figure 4. Related to Figure 2.**
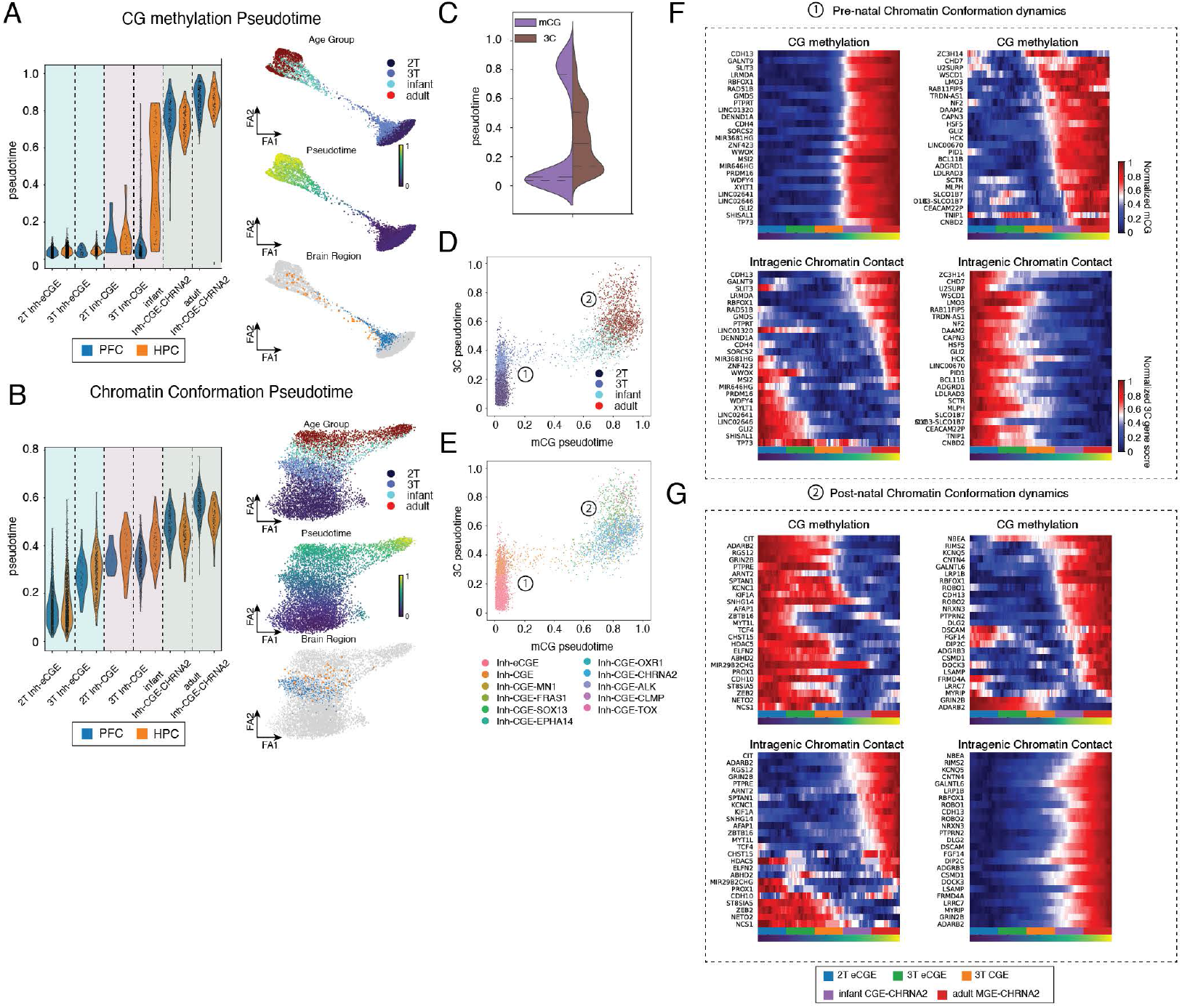
(A-B) Distribution of pseudotime scores computed from gene body mCG (A) or chromatin conformation (B) in CGE-derived inhibitory neurons. (C) Distinct distributions of pseudotime scores computed from mCG or chromatin conformation. (D-E) Direct comparison of pseudotime scores computed from mCG or chromatin conformation in individual cells, labeled by developmental age groups (D) and cell types (E). (F) mCG and 3C gene scores changes at loci associated with pre-natal chromatin conformation dynamics. (G) mCG and 3C gene scores changes at loci associated with pot-natal chromatin conformation dynamics.

**Supplementary Figure 5. Related to Figure 1-2.**
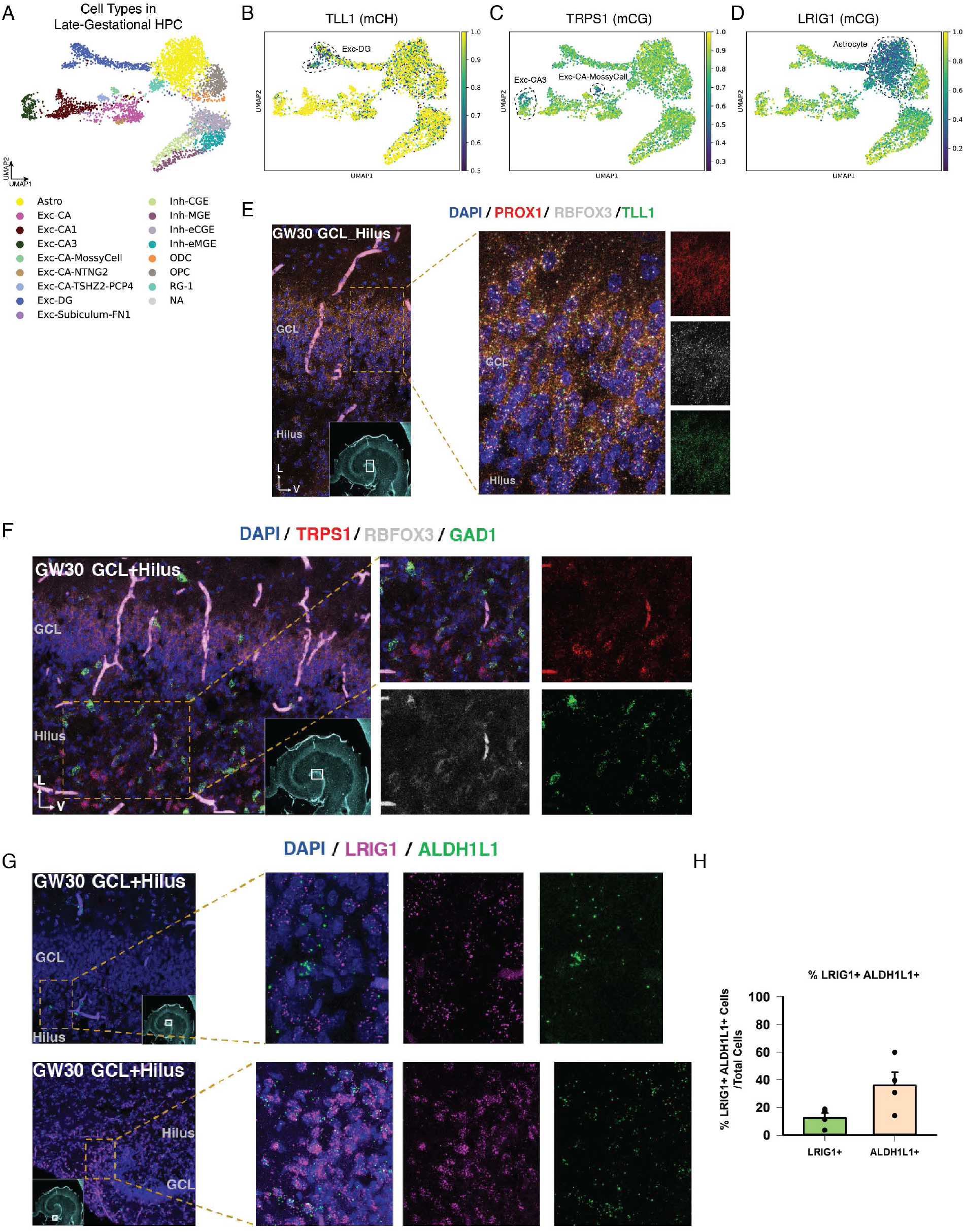
(A) UMAP of brain cells derived from late-gestational HPC samples. (B) UMAP showing TLL1 CH hypomethylation for more matured granule neurons. (C) UMAP showing TRPS1 CG hypomethylation in Mossy Cells and partially in CA3 neurons. (D) UMAP showing LRIG1 hypomethylation in astrocytes. (E) single molecular RNA in situ detection of TLL1, PROX1 and RBFOX3 transcripts in the hippocampus in the third trimester (GW 30 GW). (F) single molecular RNA in situ detection of TPRS1, GAD1 and RBFOX3 transcripts in the hippocampus in the third trimester. (G) single molecular RNA in situ detection of LRIG1 and ALDH1L1 transcripts in the hippocampus in the third trimester. (H) Quantification of LRIG1/ALDH1L1 co-expression.

**Supplementary Figure 6. Related to Figure 3.**
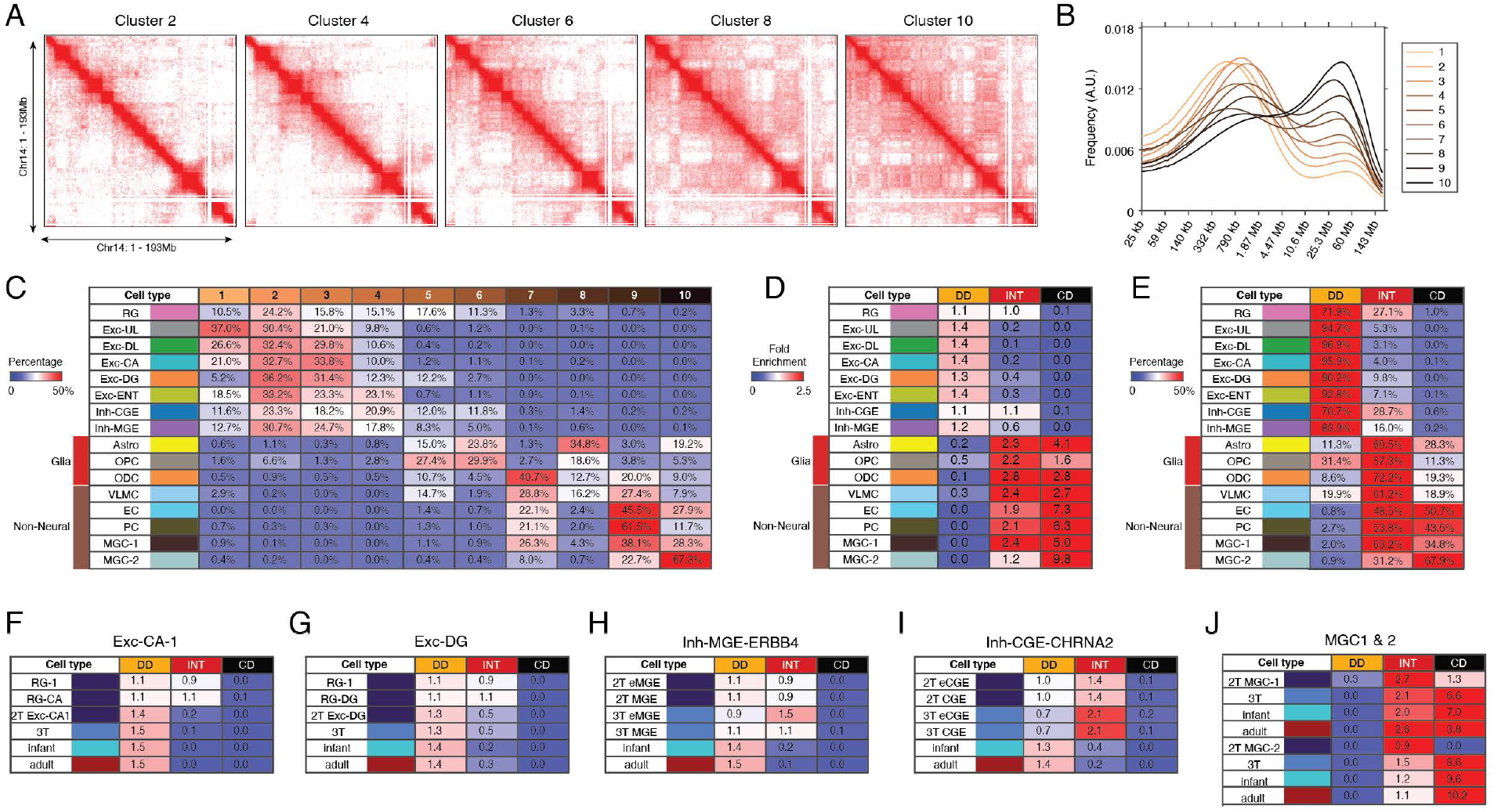
(A) Merged chromatin interaction profiles of even clusters identified in Fig. 3A. (B) Distribution of the distance between interaction loci of clusters identified in Fig. 3A. (C) Percentage of each brain cell type assigned to clusters identified in Fig. 3A. (D) Cell-type specific enrichments of DD (Domain Dominant), CD (Compartment Dominant), and INT (Intermediate) chromatin conformation. (E) Percentage of each brain cell type classified as DD, CD, or INT conformation. (F-J) Remodeling of global chromatin conformation during the differentiation of Exc-CA-1 (F), Exc-DG (G), Inh-MGE-ERBB4 (H), Inh-CGE-CHRNA2 (I), and MGC-1 & MGE-2 (J).

**Supplementary Figure 7. Related to Figure 4.**
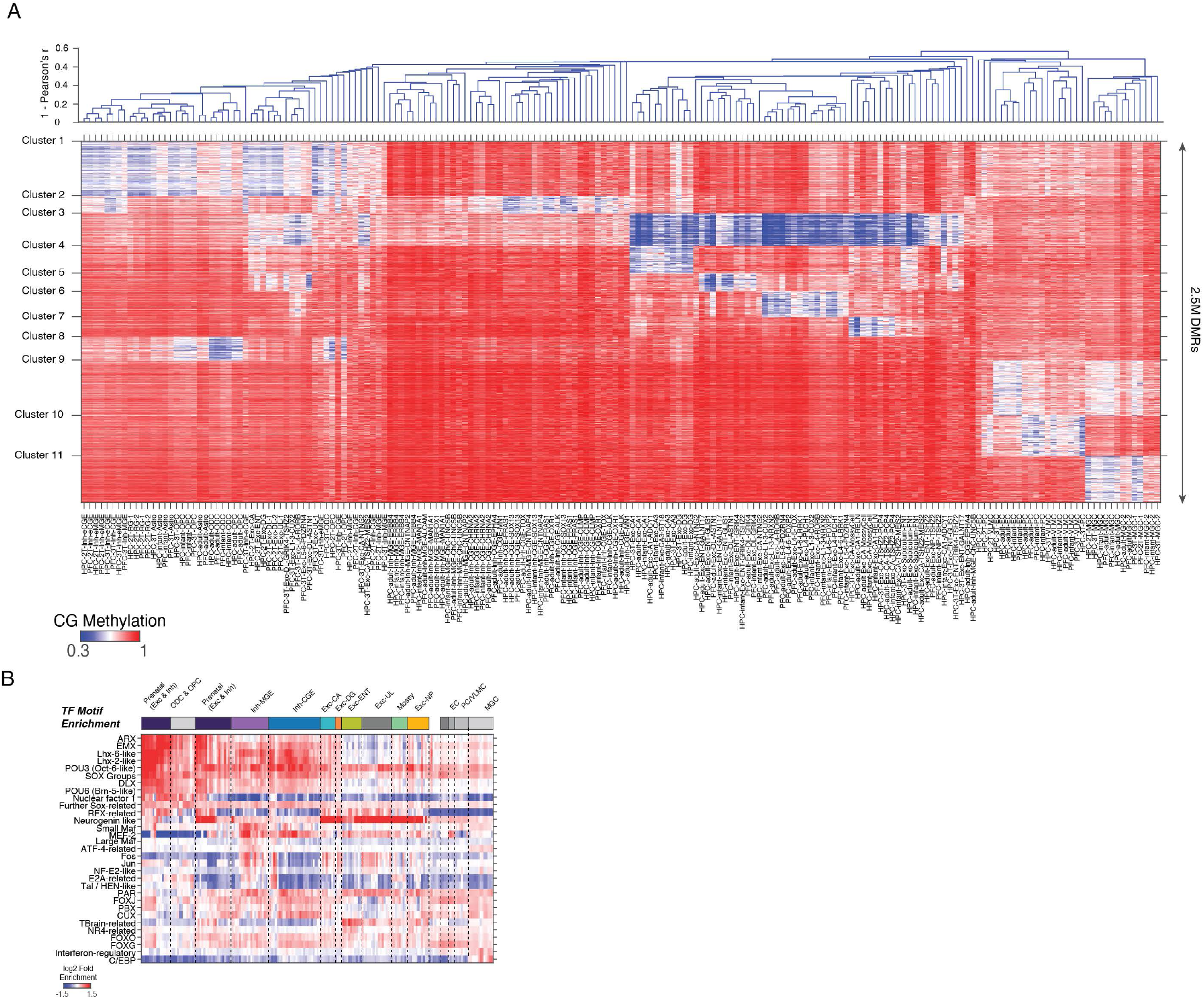
(A) K-means clustering of CG-DMRs as shown in Fig. 4A with individual cell populations labeled. (B) Transcription binding motif analysis of DMRs including statistically insignificant values (FDR> 1×10-5).

**Supplementary Figure 8. Related to Figure 4.**
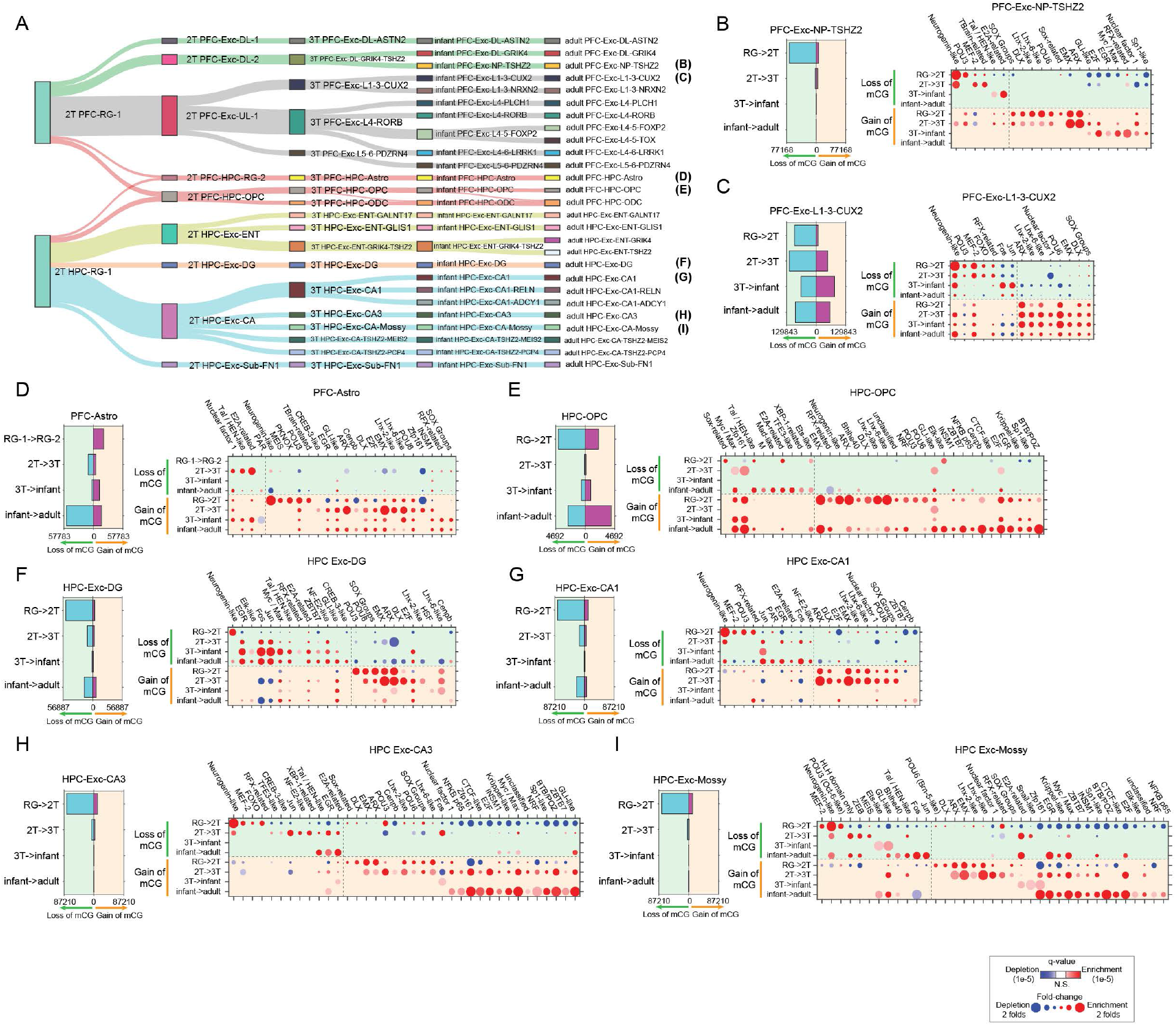
(A) Reconstructed developmental hierarchy of excitatory neurons and glial cells. (B-I) The number of trajectory-DMRs identified between developmental stages of several cell-type trajectories (left). Transcription factor binding motif enrichments between developmental stages of each trajectory. Displayed trajectories are PFC-Exc-NP-TSHZ2 (B), PFC-Exc-L1-2-CUX2 (C), PFC Astro (D), HPC-OPC (E), HPC-Exc-DG (F), HPC-Exc-CA1 (G), HPC-Exc-CA3 (H), and HPC-Exc-Mossy (I).

**Supplementary Figure 9. Related to Figure 4.**
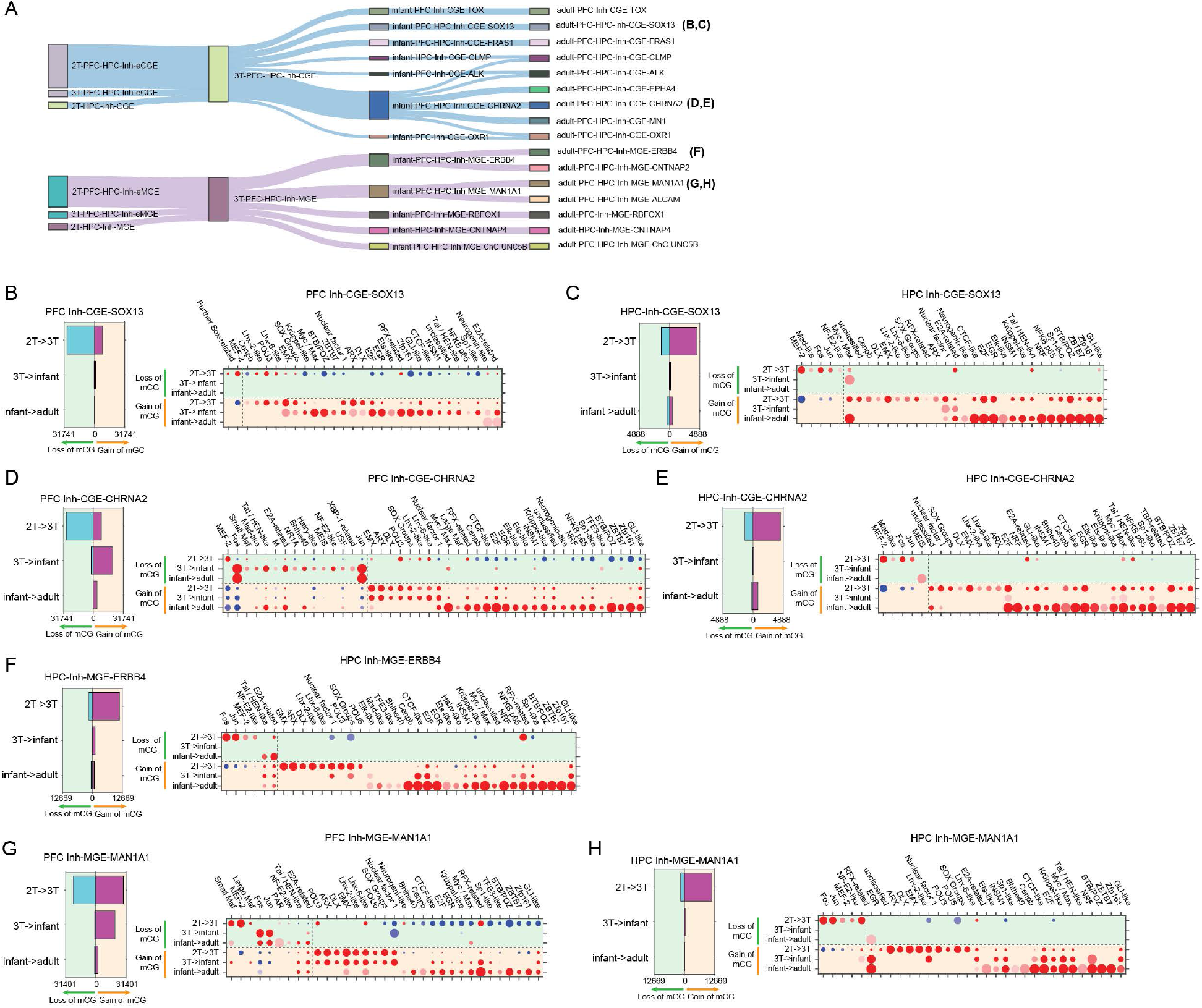
(A) Reconstructed developmental hierarchy of CGE- and MGE-derived inhibitory neurons. (B-H) Trajectory-DMRs identified between developmental stages of cell-type trajectories (left). Transcription factor binding motif enrichments between developmental stages of the trajectory. Displayed trajectories are PFC-Inh-CGE-SOX13 (B), HPC-Inh-CGE-SOX13 (C), PFC-Inh-CGE-CHRNA2 (D), HPC-Inh-CGE-CHRNA2 (E), HPC-Inh-MGE-ERBB4 (F), PFC-Inh-MGE-MAN1A1 (G), HPC-Inh-MGE-MAN1A1 (H).

**Supplementary Figure 10. Related to Figure 4.**
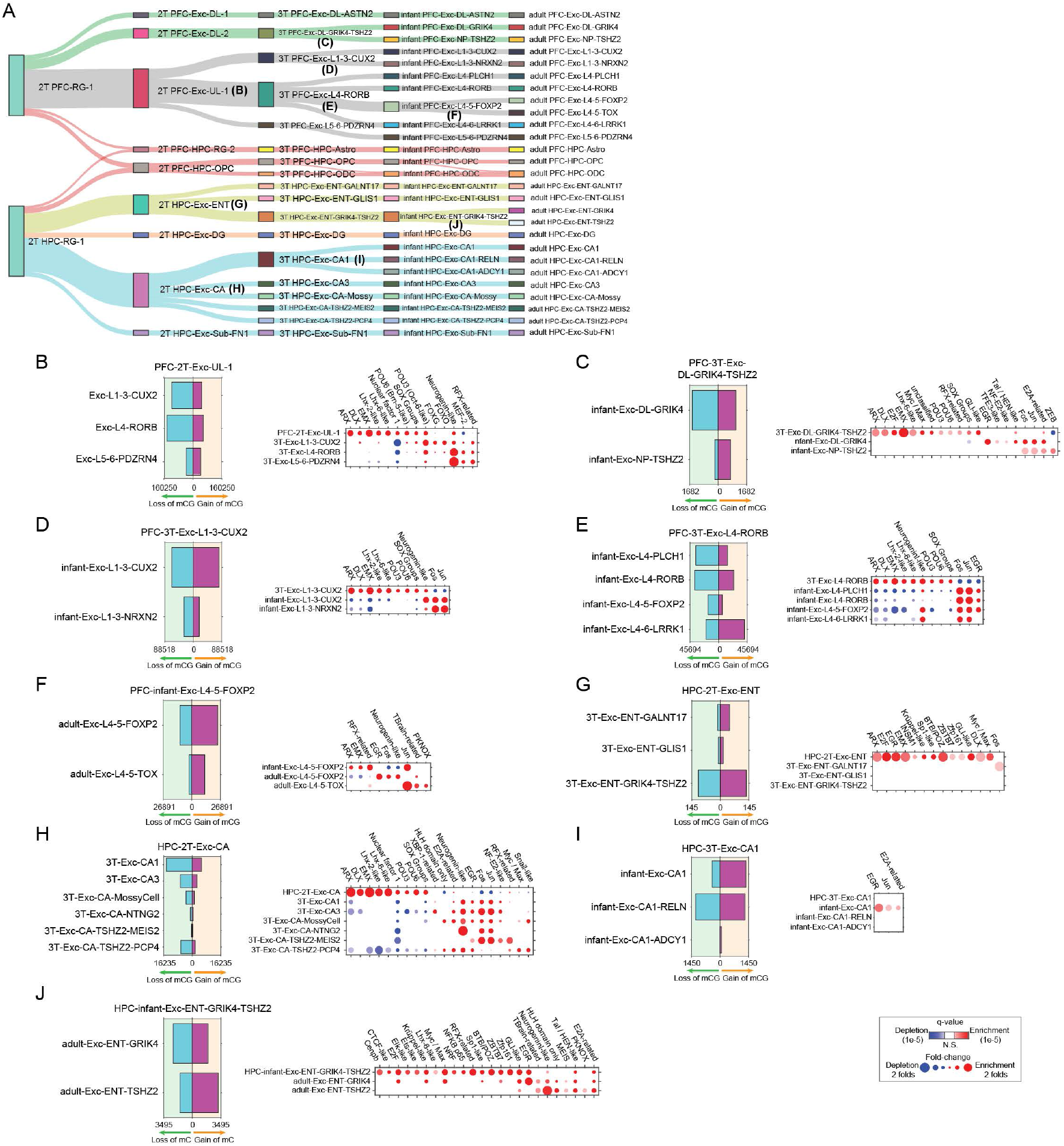
(A) Reconstructed developmental hierarchy of excitatory neurons and glial cells labeled for the branching of a mother cell type in an earlier developmental stage to daughter cell types in a later development stage. (B-J) The number of hypo-methylated branch-DMRs identified at cell-type branches and transcription factor binding motif enrichments in DMRs. Displayed cell-type branches are associated with mother cell populations PFC-2T-Exc-UL-1 (B), PFC-3T-Exc-DL-GRIK4-TSHZ2 (C), PFC-3T-Exc-L1-3-CUX2 (D), PFC-3T-Exc-L4-RORB (E), PFC-infant-Exc-L4-5-FOXP2 (F), HPC-2T-Exc-ENT (G), HPC-2T-Exc-CA (H), HPC-3T-Exc-CA1 (I) and HPC-infant-Exc-ENT-GRIK4-TSHZ2 (J).

**Supplementary Figure 11. Related to Figure 4.**
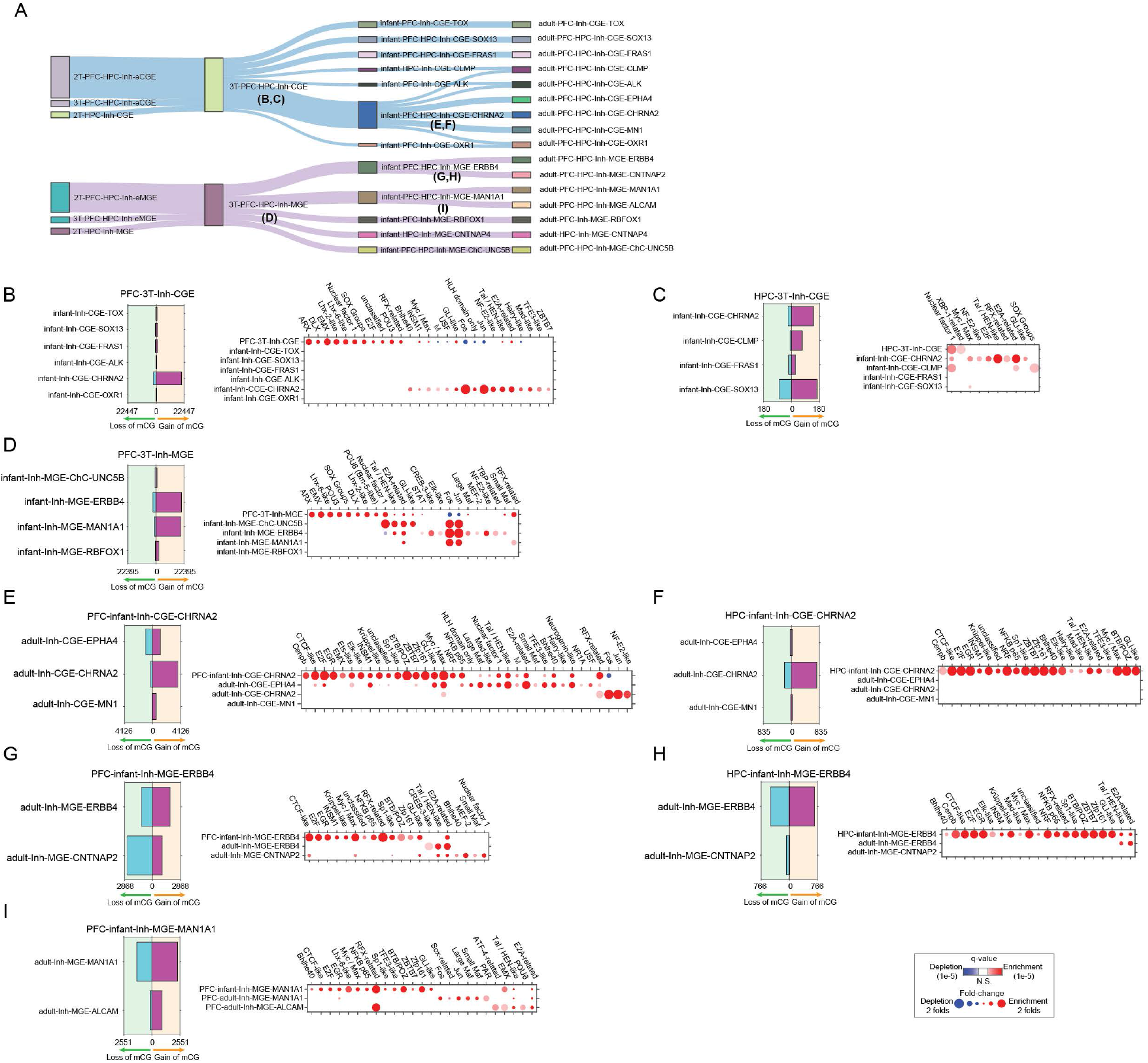
(A) Reconstructed developmental hierarchy of inhibitory neurons labeled for the branching of a mother cell type in an earlier developmental stage to daughter cell types in a later development stage. (B-J) The number of hypo-methylated branch-DMRs identified at cell-type branches and transcription factor binding motif enrichments in DMRs. Displayed cell-type branches are associated with mother cell populations PFC-3T-Inh-CGE (B), HPC-3T-Inh-CGE (C), PFC-3T-Inh-MGE (D), PFC-infant-Inh-CGE-CHRNA2 (E), HPC-infant-Inh-CGE-CHRNA2 (F), PFC-infant-Inh-MGE-ERBB4 (G), HPC-infant-Inh-MGE-ERBB4 (H), PFC-infant-Inh-MGE-MAN1A1 (I).

**Supplementary Figure 12. Related to Figure 5.**
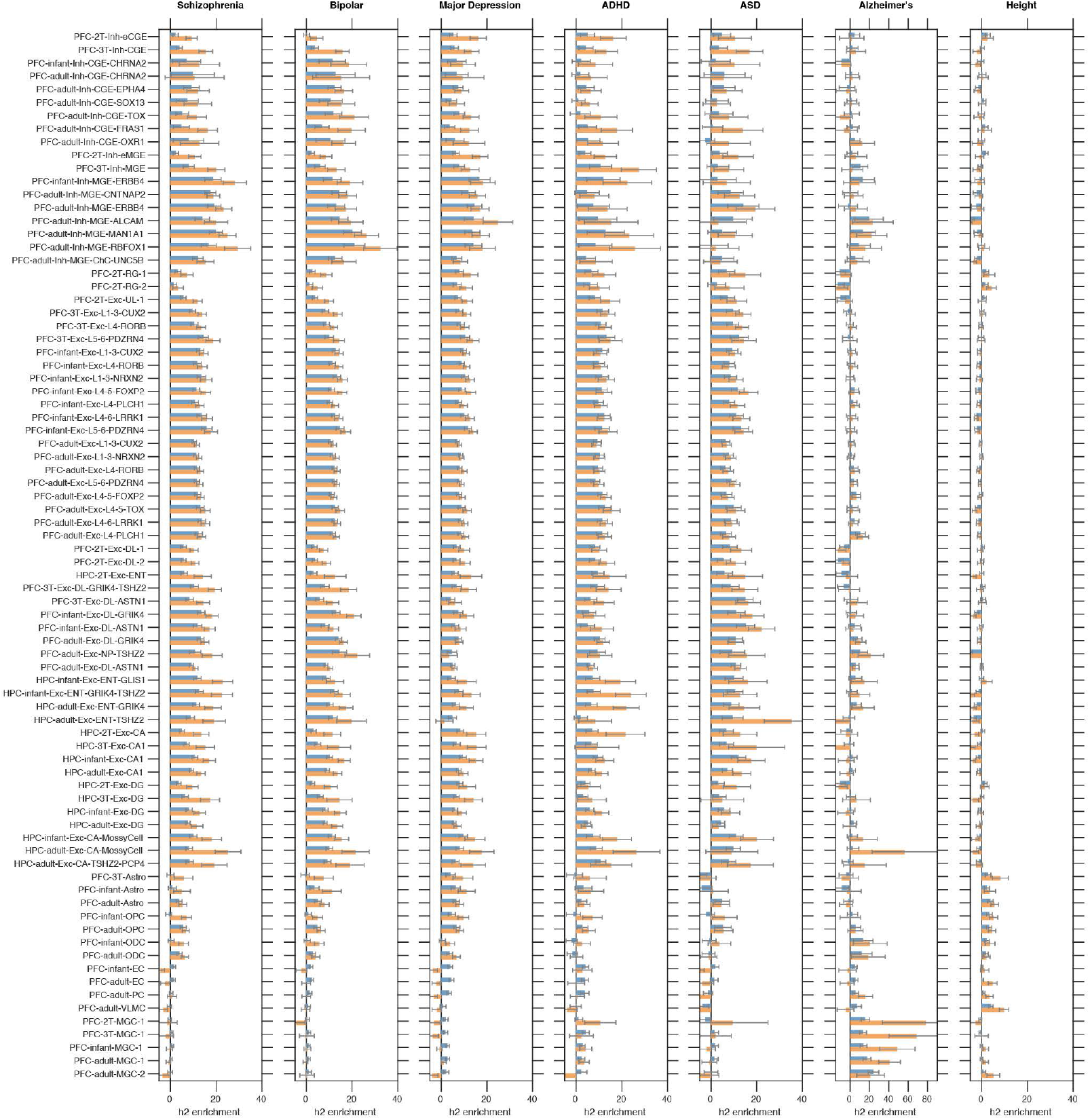
Enrichment of polygenic heritability for seven traits in DMRs (blue bars) and loop-connected DMRs (orange bars) across cell types.

**Supplementary Figure 13. Related to Figure 5.**
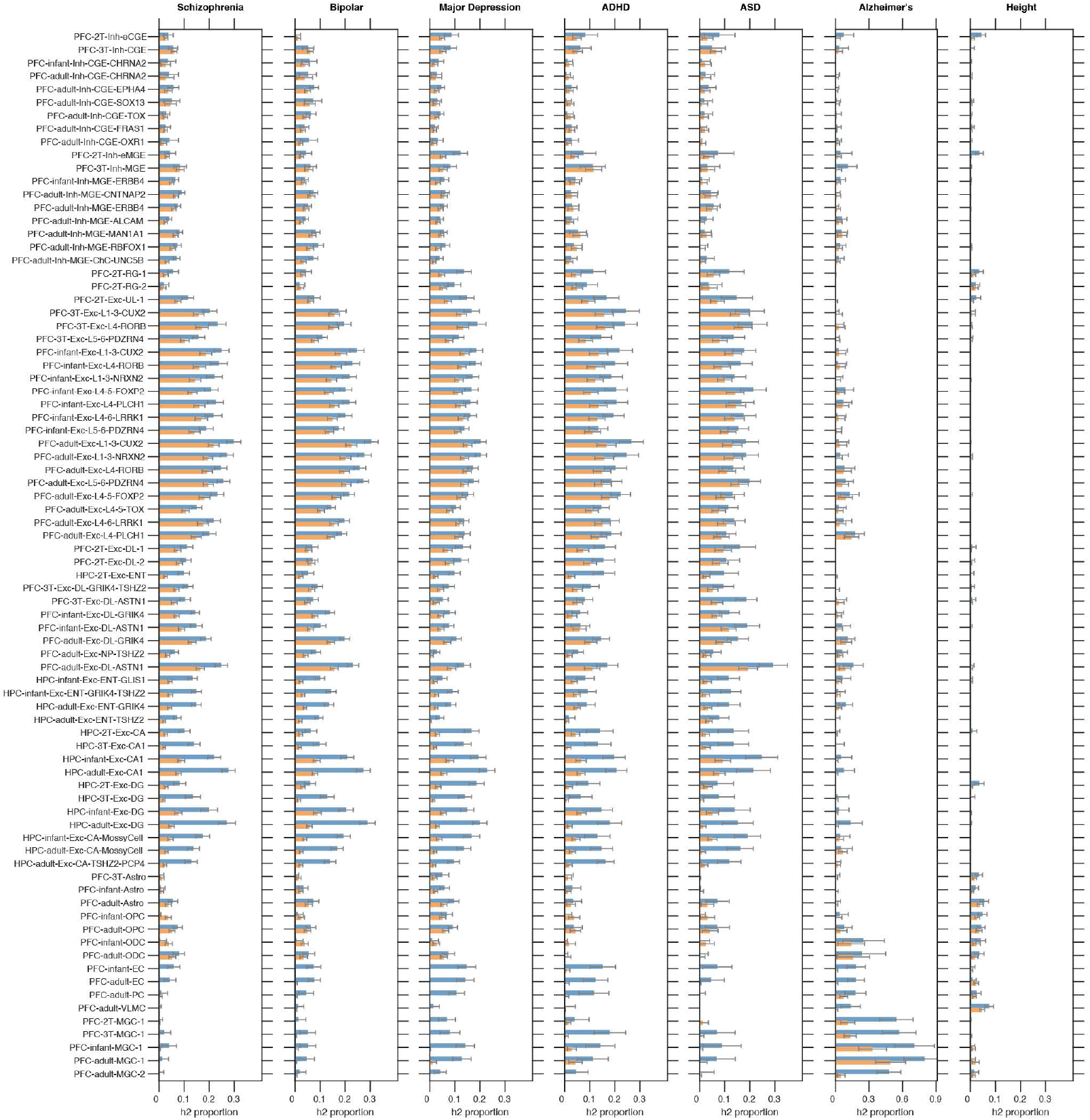
Proportion of polygenic heritability for seven traits in DMRs (blue bars) and loop-connected DMRs (orange bars) across cell types.

**Supplementary Figure 14. Related to Figure 5.**
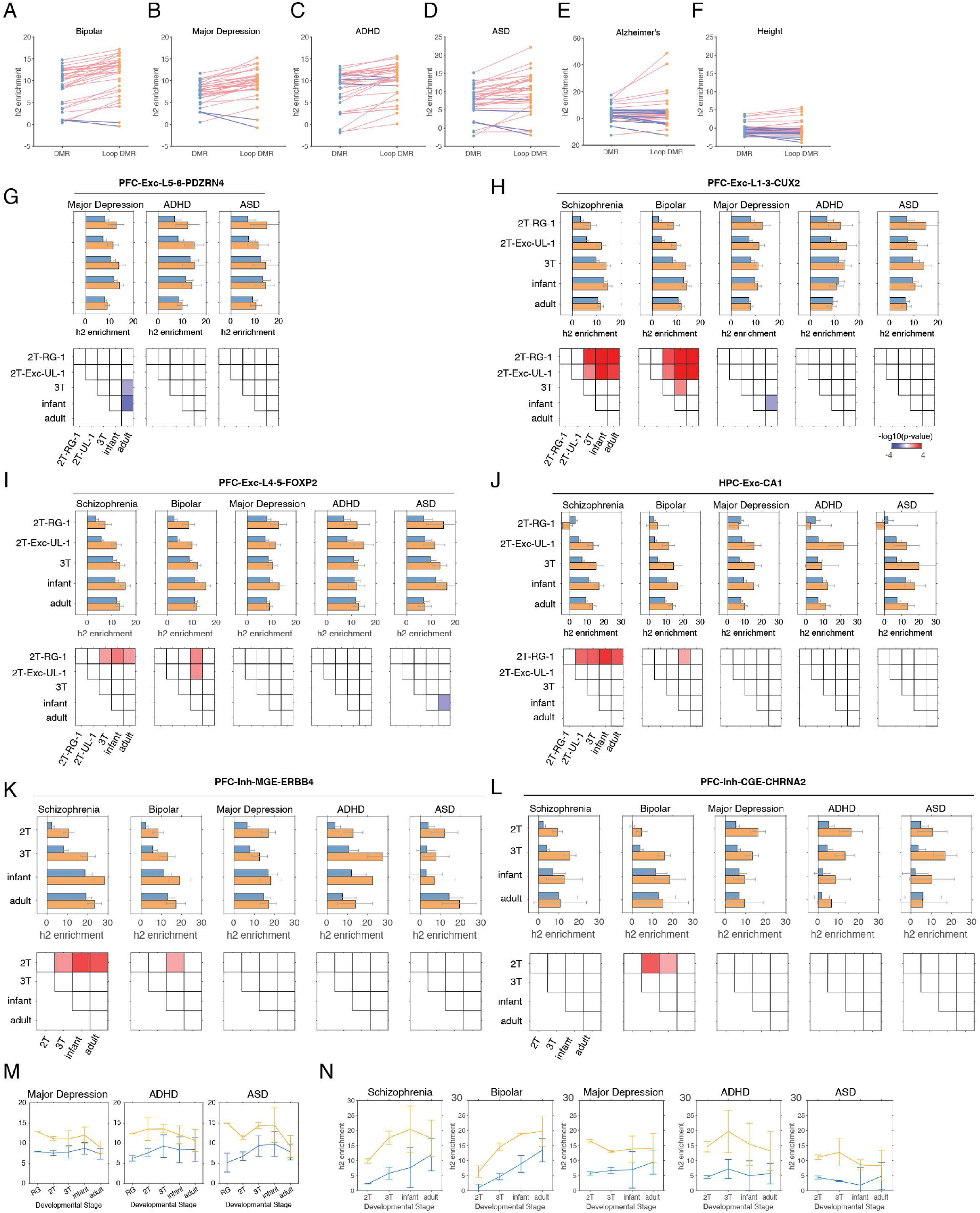
(A-F) The enrichment of polygenic heritability for bipolar disorder (A), major depression (B), ADHD (C), ASD (D), Alzheimers’ disease (E) and height (F) in DMRs and loop-connected DMRs. (G-L) Enrichment of polygenic heritability for neuropsychiatric disorders across developmental stages in PFC-Exc-L5-6-PDZRN4 (G), PFC-Exc-L1-3-CUX2 (H), PFC-Exc-L4-5-FOXP2 (I), HPC-Exc-CA1 (J), PFC-Inh-MGE-ERBB4 (K), PFC-Inh-CGE-CHRNA2 (L). (M-N) Meta-analysis of heritability enrichment for neuropsychiatric disorders in excitatory (M) and inhibitory (N) neuron populations.

## Methods

### Brain Specimen

Human Tissue Collection: Specimens were collected from autopsy with previous patient consent to institutional ethical regulations of the University of California San Francisco Committee on Human Research. Collection was at post-mortem intervals (PMI) less than 24 hours. Tissue was collected at the following institutions with previous patient consent to institutional ethical regulations of the University of California, San Francisco (UCSF) Committee on Human Research. Protocols were approved by the Human Gamete, Embryo and Stem Cell Research Committee (Institutional Review Board GESCR# 10-02693) at UCSF. Specimens were evaluated by a neuropathologist as control samples. Tissues were cut coronally and areas of interest were sampled. 1 mm tissue blocks used for chromatin and methylation assays were flash frozen in liquid nitrogen and stored in -80 C. Blocks used for histological analyses were fixed with 4% paraformaldehyde for two days, and cryoprotected in a 30% sucrose gradient. The tissue was then frozen in OCT and blocks were cut at 30 um with a cryostat and mounted onto glass slides. For each case used, we cresyl stained three sections spanning the block to ensure our position using anatomical landmarks, such as the lateral ventricle, presence of the caudate, thalamus and hippocampus.

### sn-m3C-seq

For prenatal brain samples, sn-m3C-seq was performed without the labeling of neuronal nuclei using anti-NeuN antibody, whereas post-natal samples were labeled by anti-NeuN antibody during the nuclei isolation procedure. For sn-m3C-seq performed without labeling, frozen powder of brain tissue was resuspended in 10 mL of DPBS with 2% formaldehyde and incubated at room temperature for 10 mins with slow rotation. The crosslinking reaction was quenched with 1.17 mL of 2M Glycine for 5 mins at room temperature. The crosslinked tissue sample was pellet by centrifugation with 2,000 x g for 10 mins at 4°C. The same centrifugation condition was used to pellet nuclei throughout the sn-m3C-seq procedure. The pellet was resuspended in 3mL of NIBT (10mM Tris-HCl pH=8.0, 0.25M Sucrose, 5mM MgCl_2_, 25mM KCl, 1mM DTT, 0.1% Triton X-100 and 1:100 Protease Inhibitor Cocktail (Sigma #P8340). The resuspended tissue sample was dounced with a dounce homogenizer (Sigma #D9063) for 40 times with a loose pestle and 40 times with a tight pestle. The lysate was mixed with 2 mL of 50% Iodixanol (prepared by mixing OptiPrep™ Density Gradient Medium (Sigma #D1556) with diluent (120 mM Tris-Cl pH=8.0, 150 mM KCl and 30mM MgCl_2_) with a volume ratio of 5:1). The lysate was gently layered on top of 25% Iodixanol cushion and centrifuged at 10,000 x g for 20 min at 4°C using a swing rotor. The nuclei pellet was resuspended in 1mL of cold DPBS followed by the quantification of nuclei using a Biorad TC20 Automated Cell Counter (Biorad #1450102).

In situ 3C reaction was performed using Arima Genomics Arima-HiC+ Kit. Each in situ 3C reaction uses 300K to 450K nuclei. Nuclei aliquots were pellet and resuspended in 20 µl H_2_O mixed with 24 µl Conditioning Solution and incubated at 62°C for 10 min. After the incubation, 20 µl of Stop Solution 2 was added to the reaction and incubated at 37°C for 15 min. A restriction digestion mix containing 7 µl of 10X NEB CutSmart Buffer (NEB #B7204), 4.5 µl of NlaIII (NEB #R0125), 4.5 µl of MboI (NEB #R0147) and 12 µl of 1X NEB CutSmart Buffer was added to the reaction followed by incubation at 37°C for 1 hr. The restriction digestion reaction was stopped by incubation at 65°C for 20 mins. A ligation mix containing 70 µl of Buffer C and 12 µl of Enzyme C was added and was followed by incubation at room temperature for 15 min. The reaction was then placed at 4°C overnight.

Prior to fluorescent-activated nuclei sorting (FANS), 900 µl of cold DPBS supplemented with 100 µl of Ultrapure BSA (50 mg/mL, Invitrogen #AM2618) was added to the in situ 3C reaction. To fluorescently stain nuclei, 1 µl of 1 mg/mL Hoechst 33342 was added prior to sorting. FANS was performed at UCLA Broad Stem Cell Research Center Flow Cytometry core using BD FACSAria sorters. Single nuclei were sorted into 384 well plates containing 1 µl M-Digestion Buffer containing proteinase K and ~0.05 pg Lambda DNA isolated from dcm+ E.Coli (Promega #D1501).

### Single-nucleus DNA methylome library preparation with snmC-seq3

snmC-seq3 is a modification of snmC-seq2 (*43*) that provides improved throughput and reduced cost. Key differences between snmC-seq3 and snmC-seq2 include the usage of 384 instead 8 barcoded degenerated (RP-H) primers (Table S7) for the initiation of random-primed DNA synthesis using bisulfite converted DNA as template. The expanded multiplexing allows the combining of 64 single nuclei into the downstream enzymatic reactions, which provided a 8-fold reduction of the usage of Adaptase™ and PCR reagents. In addition, the amounts of Klenow exo-, Exonuclease 1 and rSAP were reduced by 10-folds compared to snmC-seq2, providing further reduction of reagent cost.

### Processing of sn-m3C-seq data

Sequencing reads were first demultiplexed by matching the first 8 base pairs of R1 reads to the predefined well barcodes (https://github.com/luogenomics/demultiplexing). Demultiplexed reads were trimmed to remove sequencing adaptors using Cutadapt 1.18 with the following parameters in paired-end mode: -f fastq -q 20 -m 50 -a AGATCGGAAGAGCACACGTCTGAAC -A AGATCGGAAGAGCGTCGTGTAGGGA. 18 bp and 10 bp were further trimmed from the 5’- and 3’-end of R1 reads, respectively. 10 bp were trimmed from both 5’- and 3’-ends of R2 reads. sn-m3C-seq reads were mapped to hg38 reference genome using a modified Taurus-MH package (https://github.com/luogenomics/Taurus-MH) (*20*). Briefly, each read end (R1 or R2) is mapped separately using Bismark with Bowtie1 with read1 as complementary (always G to A converted) and read2 (always C to T converted) as original strand. After the first alignment, unmapped reads are retained and split into 3 pieces by 40bp, 42bp, and 40bp resulting in six subreads (read1 and read2). The subreads derived from unmapped reads were mapped separately using Bismark Bowtie1. All aligned reads are merged into BAM using Picard SortSam tool with query names sorted. For each fragment, the outermost aligned reads are chosen for the chromatin conformation map generation. The chromatin contacts with both ends mapped to the same positions were considered duplicates and removed for further analysis. Duplicates reads were removed from BAM files using Picard MarkDuplicates tool before the generation of allc files using Allcools bam-to-allc tool (https://lhqing.github.io/ALLCools/).

### Single molecule fluorescent *in situ* hybridization

smFISH was performed according to the RNAscope manual (multi-plex details). Sequences of target probes, preamplifier, amplifier, and label probe are proprietary and commercially available (Advanced Cell Diagnostics (ACD), Hayward, CA). Typically, the probes contain 20 ZZ probe pairs (approx. 50 bp/pair) covering 1000bp. Here, we used probes against human genes as single-plex probes, outlined below: Hs-MEF2C (452881), Hs-GAD1-C2 (404031-C3), Hs-RBFOX3-C2 (415591-C2), Hs-TLL1-C3 (439211), Hs-TRPS1 (831611-C3), Hs-PROX1 (530241), Hs-ALDH1L1-C3 (438881-C3), Hs-LRIG1-C2 (407421-C2).

Slides were dried at 60°C for 1 h and fixed in 4% PFA for 2 h. After several washes in PBS, slides were treated with ACD Hydrogen peroxide for 10 min and then washed in water 2x before treatment in 1x target retrieval buffer (ACD) for 5 min (at 95-100°C). After washing in water and then 100% alcohol, the slides were baked at 60ºC for 30 min. After moistening samples with water, protease treatment was performed for 15 min at 40°C in the HybEZ™ oven. Hybridization of probes and amplification was performed according to the manufacturer’s instructions. In short, tissue sections were incubated in desired probe (2–3 drops/section) for 2 h at 40°C in the HybEZ™ oven. The slides were washed twice in 1x wash buffer (ACD) for 2 min each and incubated in 5X SSC at RT overnight. Amplification and detection steps were performed using the Multiplex kit (ACD, 320293) for single-plex probes. The following was performed in repeated cycle for each probe. ~4 DROPS of AMP x-FL was added to entirely cover each section and slide placed in the HybEZ™ Oven. The slide was incubated for 30 MIN at 40°C. Slides were removed from HybEZ™ Slide Rack and excess liquid removed before being submerged in the Tissue-Tek® Staining Dish filled with 1X WASH BUFFER. Slides were washed in 1X Wash Buffer for 2 MIN at RT. The next AMP x-FL was added and the cycle was repeated. Slides were washed in PBST, incubated with DAPI for 30 sec at RT and mounted in Aqua Mount (Lerner). Images were taken using 100x objective on the Leica Stellaris confocal microscope.

### Single-cell bimodal data quality control and preprocessing

Cells were filtered on the basis of several metadata metrics: (1) mCCC level <0.03; (2) global mCG level >0.5; (3) global mCH level < 0.2; and (4) Total 3C interactions >100,000. Methylation features were calculated as fractions of methylcytosine over total cytosine across gene bodies ± 2kb flanking regions and 100kb bins spanning the entire genome. Methylation features were further split into CG and CH methylation types. These features were then filtered on mean coverage s10 and values with coverage <5 were imputed as the mean feature value by sample. Principal component analysis was then run using Scanpy(*44*) default parameters followed by k-nearest neighbors (knn) using only the top 20 principal components by the amount of variance explained and k=15. Iterative clustering was then performed with a combination of leiden unsupervised clustering and UMAP dimensionality reduction, identifying clusters as cell types by marker gene hypomethylation. We observe certain batch effects in our dataset that are associated with the time the data was generated. Harmony (*45*) is used on metadata features to mitigate batch occurring between samples in the principal component feature space.

### Pseudotime Analysis

Pseudotime analysis was run following the methods outlined in (Wolf et al. 2019)(*23*). Each pseudotime analysis had clustering preprocessing steps, PCA, knn with k=15 using 20 PCs, and leiden, recomputed for its respective subset of the data.The computed leiden clusters were then used to initialize a partition-based graph abstraction (PAGA). This PAGA is used as the precomputed initialization coordinates for the visualization with force-directed graph drawing by the ForceAtlas2 package. (*46*) A root node is then set in the leiden cluster furthest from the adult cell types and Scanpy’s implementation of diffusion-based pseudotime was used. Genes are selected for display compared to the pseudotime scores by sorting by correlation and anticorrelation to the pseudotime score as well as requiring the 3C gene score to have variance >.1. For Fig. 2H and Fig. S4F, gene examples were selected by highest gene body mCG correlation to the pseudotime and 3C gene score anticorrelation. For Fig. 2I and Fig. S4G, gene examples were selected by highest gene body mCG anticorrelation to the pseudotime and 3C gene score correlation to the pseudotime. Distribution comparisons are computed by the Wilcoxon rank-sum test.

### DMR and Transcription Factor (TF) Binding Motif Analysis

All CG-DMRs were identified from pseudobulk allc files using Methylpy (https://github.com/yupenghe/methylpy) (*47*). DMRs identified from a multi-sample comparison of all cell types were used for analyses in Fig. 4A-B and Fig S7, as well as disease heritability enrichment analyses shown in Fig. 5 and Fig. S12-14. Trajectory-DMRs were identified using pairwise comparisons of adjacent development stages of a cell-type trajectory. Branch-DMRs were identified using multi-sample comparisons, including the mother cell population from an earlier developmental stage, and daughter populations from a later developmental stage. TF binding motif enrichment analysis was performed similarly as previously described (*13, 25, 48*). DMR regions were liftovered to hg19 reference genome for the TF binding motif enrichment analysis. TF binding position weight matrices (PWM) were obtained from the MEME motif database and scanned across the human hg19 reference genome to identify hits using FIMO (-- output-pthresh 1E-5, -- max-stored-scores 500000 and --max-strand) (*49, 50*). DMRs were extended 250 bp both upstream and downstream for overlapping with TF binding motif hits. The overlap between TF binding motif hits and DMRs (extended ±250 bp) were determined requiring a minimal of 1 bp overlap. The enrichment of TF binding motifs in DMSs was assessed using DMRs (extended 250 bp from center) identified across adult human tissues (tissue DMRs) as the background (*47*). The overlaps between TF binding motif hits and the foreground DMR list was compared to the overlaps between TF binding motifs hits and tissue DMRs (background) using the hypergeometric test (MATLAB hygecdf).

### Single-cell Embedding Based on Chromatin Contact

Single-cell contact matrices at 100 kb resolution were imputed by scHiCluster (*51*) with pad = 1. The imputed contacts with distance >=100 kb and <=1 Mb were used as features for singular value decomposition (SVD) dimension reduction. Principal components were normalized by singular values and L2 norms per cell, and then used for k-NN graph construction (k=25) and UMAP. 25 dimensions were used for the full dataset (Fig. 1, S2B, E), 20 dimensions were used for the RG subtypes (Fig. S2F), and 10 dimensions were used for the MGE or astrocyte lineage (Fig. S2D, H).

### Chromatin Loop and Differential Loop Analysis

Chromatin loops were identified with scHiCluster (*51*) for every cell type identified in this study. To identify loops from a group of cells, single cell contact matrices at 10kb resolution were imputed by scHiCluster with pad = 2 for the contacts within 5.05 Mb (result denoted as Q_cell_). We only performed loop calling between 50 kb and 5 Mb, given that increasing the distance only leads to a limited increase on the number of significant loops. For each single cell, the imputed matrix of each chromosome was log-transformed, and Z-score normalized at each diagonal (result denoted as E_cell_) and subtract a local background between >=30kb and <=50kb (result denoted as T_cell_), similar to SnapHiC (*52*). A pseudobulk level t-statistic was computed to quantify the deviation of E and T from 0 across single cells from the cell group, where larger deviations represent higher enrichment against global (E) or local (T) background. E_cell_ is also shuffled across each diagonal to generate E_shufflecell_, and then T_shufflecell_, to estimate a background of the t-statistics. An empirical FDR can be derived by comparing the t-statistics of observed cells versus shuffled cells. We required the pixels to have average E >0, fold change >1.33 against donut and bottom left backgrounds, fold change >1.2 against horizontal and vertical backgrounds (*52*), and FDR <0.01 compared to global and local background.

Differential loops were identified between age groups within the same major lineage. To compare the interaction strength of loops between different groups of cells, we adopt an analysis of variance (ANOVA) framework to compute the F statistics for each loop identified in at least one cell group using either Q_cell_ (result denoted as F_Q_) or T_cell_ (result denoted as F_T_). We log-transformed and then Z-scored F_Q_ and F_T_ across all the loops being tested and selected the ones with both F_Q_ and F_T_ > 1.036 (85th percentile of standard normal distribution) as differential loops. The threshold was decided by visually inspecting the contact maps as well as the correlation of interaction and loop anchor CG methylation.

### Identification of domains and differential domain boundaries

Single cell contact matrices at 25kb resolution were imputed by scHiCluster (*51*) with pad = 2 for the contacts within 10.05 Mb. Domains were identified within each single cell. Insulation scores were computed in each cell group (major type or major type within a brain region) for each bin with the pseudobulk imputed matrices (average over single cells) and a window size of 10 bins. Boundary probability of a bin is defined as the proportion of cells having the bin called as a domain boundary among the total number of cells from the group.

To identify differential domain boundaries between n cell groups, we derived an nx2 contingency table for each 25kb bin, where the values in each row represent the number of cells from the group that has the bin called as a boundary or not as a boundary. We computed the Chi-square statistic and p-value of each bin, and used the peaks of the statistics across the genome as differential boundaries. The peaks are defined as local maximum of Chi-square statistics, within FDR <1e-3 (Benjamini and Hochberg procedure). If two peaks are within 5 bins to each other, we only kept the peak with a higher Chi-Square statistic. We also require the peaks to have a Z-score transformed Chi-square statistic >1.960 (97.5 percentile of standard normal distribution), fold-changes between maximum and minimum insulation score >1.2, and differences between maximum and minimum boundary probability >0.05.

### 3C Gene Score

3C gene score is defined as the sum of off-diagonal values from the row and column of the TSS bin to the row and column of the TES bin in the imputed 10kb contact matrices.

### Chromatin Potential Analysis

We considered the methylation and 3C profile of a cell separately as a 3C cell and a mC cell, then projected the mC cells and 3C cells to the same low-dimensional space based on their molecular similarities across 100kb genome bins. This is achieved by a 3-step method analogous to Seurat v3 (*53*): 1) Using canonical correlation analysis (CCA) to capture the shared variance between 3C cells and mC cells; 2) finding anchors as 5 mutual nearest neighbors (MNN) between the two modalities; 3) pulling the two modalities into the same space. Specifically, we started from all the 100kb-bin-pairs with both anchors passed methylation read coverage filters, and used ALLCools to select group (cell type by age group) enriched features. For mC cells, we used the average mCG level of the two anchors after normalizing the global mCG level of each cell. For 3C cells, we used the imputed contact values at 100kb resolution. CCA was performed as *USV*′ = Σ_*c*_ *XcYc*, where *Xc* and *Yc* are cell-by-binpair matrices representing the feature matrix of chromosome c after Z-score normalization of each feature across cells. U and V were normalized by dividing the L2-norm of each row, and used to find MNN anchors and score anchors using the same method as Seurat v3. *Xc* and *Yc* were also combined vertically and the PCs of this combined matrix were integrated together using the same method as Seurat v3 through the anchors generated from the previous step. This integration step projects the PCs of 3C cells to mC cells while keeping the PCs of mC cells unchanged.

After joint embedding of mC cells and 3C cells, for each 3C cell (query), we find its nearest 5 mC cells on the embedding space, representing the cells whose methylation profiles are the most similar to the 3C profile of the query cell, and thus define a “3D chromatin potential” by an arrow pointing from the 3C cell to the average of its mC neighbors. The discrepancy between the real corresponding mC-3C profile (from the same cell) and the inferred correspondence suggest the timing differences between the two modalities during development. For instance, an arrow pointing from mid-gestation to late-gestation represents that the 3C profile of the mid-gestation cell is more correlated with the methylation profile of late-gestation cells than the methylation profile of itself, suggesting 3C changes earlier than methylation. It is worth noting that the result shows a general pattern between the two modalities across the genome, while the dynamics at individual genes could be different, which needs to be studied specifically.

### Distribution of the distance between interacting loci analysis

In order to count the number of cis (intra-chromosomal) contacts in each cell and bulk Hi-C data (*28*), we divided the contacts into 143 logarithmic bins, the first of which was for contacts that were separated by less than 1 Kb. Each subsequent bin covered an exponent step of 0.125, using base 2. Contacts in bins 1-37 were determined to be noisy and were eliminated, leaving bins 38-141 as the valid bins.

The following metrics were used for the following analysis.

- % near - percentage of contacts in bins 38-89 out of all valid bins
- % long - percentage of contacts in bins 90-141 out of all valid bins

Cells were clustered by the distribution of their distance between interacting loci (k-means, k = 10) and reordered by the average value of log2(% near/% long) of each cluster (Fig. 3A-D and Fig. S6B-C).

Each cell was assigned to a group by following criteria (Fig. 3E-F and Fig. S6D-J):

- Domain dominant (DD): log2(% near/% long) >=0.4
- Intermediate (INT): -0.4 > log2(% near/% long) >0.4
- Compartment dominant (CD): log2(% near/% long) <=-0.4

To find the cluster enriched in each cell type, we first calculated the percentage belonging to each cluster by cell type (Fig. S6C,E). The enrichment score was obtained by normalizing the fraction of each cell type by the relative cluster sizes (Fig. 3C,E,F and Fig. S6D,F-J)).

### Polygenic heritability enrichment analysis

Polygenic heritability enrichment of DMR and/or chromatin loop was performed using S-LDSC partitioned heritability analysis (*54*). GWAS summary statistics included Schizophrenia (*36*), Bipolar’s disorder (*55*), Major Depressive Disorder (*56*), ADHD (*57*), ASD (*58*), Alzheimer’s disease (*59*), Height GWAS in UK Biobank (*60*) (downloaded from https://alkesgroup.broadinstitute.org/sumstats_formatted/). For each cell type, binary annotations were created using DMR and/or chromatin loop. We considered two types of genomic regions: (1) DMR: including all DMRs for a given cell type. (2) loop-connected DMR: including the subset of DMRs that overlap with any of the chromatin loop called in the matching cell types. To create binary annotations, SNPs in these genomics regions were assigned as 1 and otherwise 0. Then we assessed the heritability enrichment of each of these annotations conditional on the “baseline model” (*35*). We reported heritability enrichment and proportion of heritability using ‘Enrichment’, ‘Enrichment_std_error’, ‘Prop._h2’, ‘Prop._h2_std_error’ columns in S-LDSC results. To assess statistical significance for heritability enrichment differences across annotations (e.g., differences between cell types in a developmental trajectory), we used t-test to test the differences of heritability enrichment of two cell types with d.o.f. = 200 + 200 - 2, where 200 corresponds to the number of jackknife samples in S-LDSC block jackknife procedure.

### Overlap between fine-mapped variants and DMR / chromatin loop for schizophrenia

We used statistical fine-mapping results that were previously performed in the latest PGC schizophrenia study (*36*). We filtered for autosomal high-confidence putative causal SNPs with PIP > 10%, and retained 190 independent association loci (containing 569 SNPs in total), with each loci containing a credible set with 3.0 SNPs on average. We used Fisher’s exact test to assess the overlap between these 569 fine-mapped SNPs and DMR/chromatin loop annotations using all SNPs in GWAS summary statistics as background (see above for constructing DMR/chromatin loop annotations). We reported odds ratios of the overlap. We also assessed the overlap between 190 schizophrenia fine-mapped loci (as aggregates of 569 putative causal SNPs) and DMR/chromatin loop annotations (Fig. 5B). We define the overlap between fine-mapped loci and DMR/chromatin loop annotations based on whether any high-confidence putative causal SNP in the fine-mapped loci located in the annotation. Furthermore, we overlapped putative causal SNP and DMR/chromatin loop annotations to GTEx high-confidence fine-mapped cis-eQTL data (downloaded from https://storage.googleapis.com/gtex_analysis_v8/single_tissue_qtl_data/GTEx_v8_finemapping_CAVIAR.tar): we first identified SNP-gene pairs such that the putative causal SNP is located in DMRs and connected to transcription start site of any gene via chromatin loops, and then we overlapped these SNP-gene pairs with eQTL cis-eQTL/eGene pairs.

## Notes

https://brain-epigenome.cells.ucsc.edu/

